# Regional differences in synaptic degeneration are linked to alpha-synuclein burden and axonal damage in Parkinson’s disease and Dementia with Lewy bodies

**DOI:** 10.1101/2023.12.07.570569

**Authors:** Irene Frigerio, Maud MA Bouwman, Ruby TGMM Noordermeer, Ema Podobnik, Marko Popovic, Evelien Timmermans, Annemieke JM Rozemuller, Wilma DJ van de Berg, Laura E Jonkman

**Author notes:** Correspondence to: Irene Frigerio De Boelelaan 1118, 1081 HV, Amsterdam, the Netherlands.

## Abstract

Regional differences in synaptic degeneration may underlie differences in clinical presentation and neuropathological disease progression in Parkinson’s Disease (PD) and Dementia with Lewy bodies (DLB). Here, we mapped and quantified synaptic degeneration in cortical brain regions in PD, PD with dementia (PDD) and DLB, and assessed whether regional differences in synaptic loss are linked to axonal degeneration and neuropathological burden. We included a total of 47 brain donors, 9 PD, 12 PDD, 6 DLB and 20 non-neurological controls. Synaptophysin^+^ and SV2A^+^ puncta were quantified in eight cortical regions using a high throughput microscopy approach. Neurofilament light chain (NfL) immunoreactivity, Lewy body (LB) density, phosphorylated-tau and amyloid-β load were also quantified. Group differences in synaptic density, and associations with neuropathological markers and Clinical Dementia Rating (CDR) scores, were investigated using linear mixed models. We found significantly decreased synaptophysin and SV2A densities in the cortex of PD, PDD and DLB cases compared to controls. Specifically, synaptic density was decreased in cortical regions affected at Braak α-synuclein stage 5 in PD (middle temporal gyrus, anterior cingulate and insula), and was additionally decreased in cortical regions affected at Braak α-synuclein stage 4 in PDD and DLB compared to controls (entorhinal cortex, parahippocampal gyrus and fusiform gyrus). Synaptic loss associated with higher NfL immunoreactivity and LB density. Global synaptophysin loss associated with longer disease duration and higher CDR scores. Synaptic neurodegeneration occurred in temporal, cingulate and insular cortices in PD, as well as in parahippocampal regions in PDD and DLB. In addition, synaptic loss was linked to axonal damage and severe α-synuclein burden. These results, together with the association between synaptic loss and disease progression and cognitive impairment, indicate that regional synaptic loss may underlie clinical differences between PD and PDD/DLB. Our results might provide useful information for the interpretation of synaptic biomarkers *in vivo*.

## Introduction

Synaptic loss is an early phenomena in Parkinson’s disease (PD) and Dementia with Lewy bodies (DLB) [1, 2]. PD is the second most common neurodegenerative disorder, affecting almost 3% of the population over 65 years old [3], and 80% of PD patients develop dementia (PDD) after 20 years of disease onset [4–7]. When patients develop dementia before or within one year of motor symptom onset, the disease is diagnosed as DLB [8]. Pathologically, PD(D) and DLB are synucleinopathies [9], with neuropathological hallmarks of Lewy bodies (LBs) and Lewy neurites [10], consisting mainly of the protein α-synuclein, disrupted organelles, membranes and lipids [11]. In physiological conditions, α-synuclein is abundant in the pre-synaptic compartment [12]. Post-mortem and experimental studies provided evidence that α-synuclein aggregation might start at pre-synaptic terminals, inducing synaptic degeneration and subsequently neuronal loss [1, 13, 14]. In post-mortem brains, a loss of synaptophysin [15] and several other synaptic proteins [16] was reported in frontal, occipital and hippocampal regions in PD, and in the parietal cortex in DLB [17]. However, there is still limited knowledge on the regional patterns of synaptic loss in PD, PDD and DLB, as these studies only included a limited number of regions.

Since PD is increasingly recognized as a synaptopathy [13], many efforts have been made to identify fluid biomarkers of synaptic loss [18–21], and imaging positron emission tomography (PET) tracers reflecting synaptic density [22]. The increasing efforts have led to the discovery that in PD and DLB patients, levels of several synaptic proteins are altered in CSF [18–21], and that binding of the PET tracer synaptic vesicle glycoprotein 2 (SV2A) is reduced in the brainstem and cortex [23–25], even though regional results remain inconsistent across studies. Moreover, limited information is available on the neuropathological alterations linked to synaptic loss in PD, PDD and DLB. In fact, besides α-synuclein pathology, Alzheimer’s disease (AD) co-pathology [26–28] and axonal degeneration [29, 30] are often present in the cortex of PDD and DLB patients, and might contribute to regional synaptic degeneration in these phenotypes. Taken together, studying regional differences in synaptic degeneration, and which neuropathological processes contribute to it, may provide insights into the selective vulnerability and neuropathological disease progression of PD(D) and DLB cases.

As such, the aim of this study was to map and quantify synaptic loss with the pre-synaptic markers synaptophysin and SV2A in eight cortical brain regions commonly affected in PD(D) and DLB, and to assess whether regional differences in the pattern of synaptic loss are linked to axonal degeneration and neuropathological burden. We hypothesized that cortical brain regions affected at Braak α-synuclein stage 4, such as the entorhinal cortex, parahippocampal and fusiform gyrus, show more pronounced synaptic loss than regions affected at stages 5 and 6, such as the temporal, cingulate, insular, and frontal cortices. Moreover, we hypothesized that axonal degeneration and neuropathological burden, especially α-synuclein load, correlate with synaptic loss. The results of this study might provide useful information for the interpretation of synaptic biomarkers *in vivo*.

## Materials and methods

### Donor inclusion

In collaboration with the Netherlands Brain Bank (NBB; http://brainbank.nl) and Normal Aging Brain Collection Amsterdam (NABCA; http://nabca.eu) [31] we included a total of 47 brain donors. 12 PD, 9 PDD and 6 DLB were included based on clinical presentation [7, 8, 32]. Age at diagnosis (symptoms onset), disease duration (age at death minus age at diagnosis), motor-to-dementia interval (years between motor symptoms and dementia onset) and Clinical Dementia Rating scores (CDR) [33] were extracted from the donor’s clinical files. Neuropathological diagnosis was confirmed by an expert neuropathologist (A.R.) and performed according to the international guidelines of the Brain Net Europe II (BNE) consortium (https://www.brainbank.nl/about-us/brain-net-europe/) [34–36]. Additionally, 20 non-neurological controls were included from the NABCA database [31]. All donors signed an informed consent for brain donation and the use of material and clinical information for research purposes. The procedures for brain tissue collection of NBB and NABCA have been approved by the Medical Ethical Committee of Amsterdam UMC, VUmc. For donor characteristics, see **Table S1**.

### Tissue sampling

Workflow of the methods is visualized in **Figure 1**. Brain tissue was collected at autopsy, resulting in an average post-mortem delay (interval between death and autopsy) of 8 hours. Eight formalin-fixed paraffin-embedded tissue blocks (4%, four weeks fixation) were obtained from the right hemisphere and processed for immunohistochemistry. Based on Braak α-synuclein staging [10], we defined 3 groups of brain regions: (i) regions affected at Braak α-synuclein stage 4, consisting of the entorhinal cortex (ENTC), parahippocampal gyrus (PHG) and fusiform gyrus (FusG); (ii) regions affected at Braak α-synuclein stage 5, consisting of the anterior cingulate gyrus (ACC), anterior insula (AI), and middle temporal gyrus (MTG); (iii) regions affected at Braak α-synuclein stage 6, consisting of the superior frontal gyrus (GFS) and the posterior cingulate gyrus (PCC; the parietal cortex was not available).

**Fig. 1.**
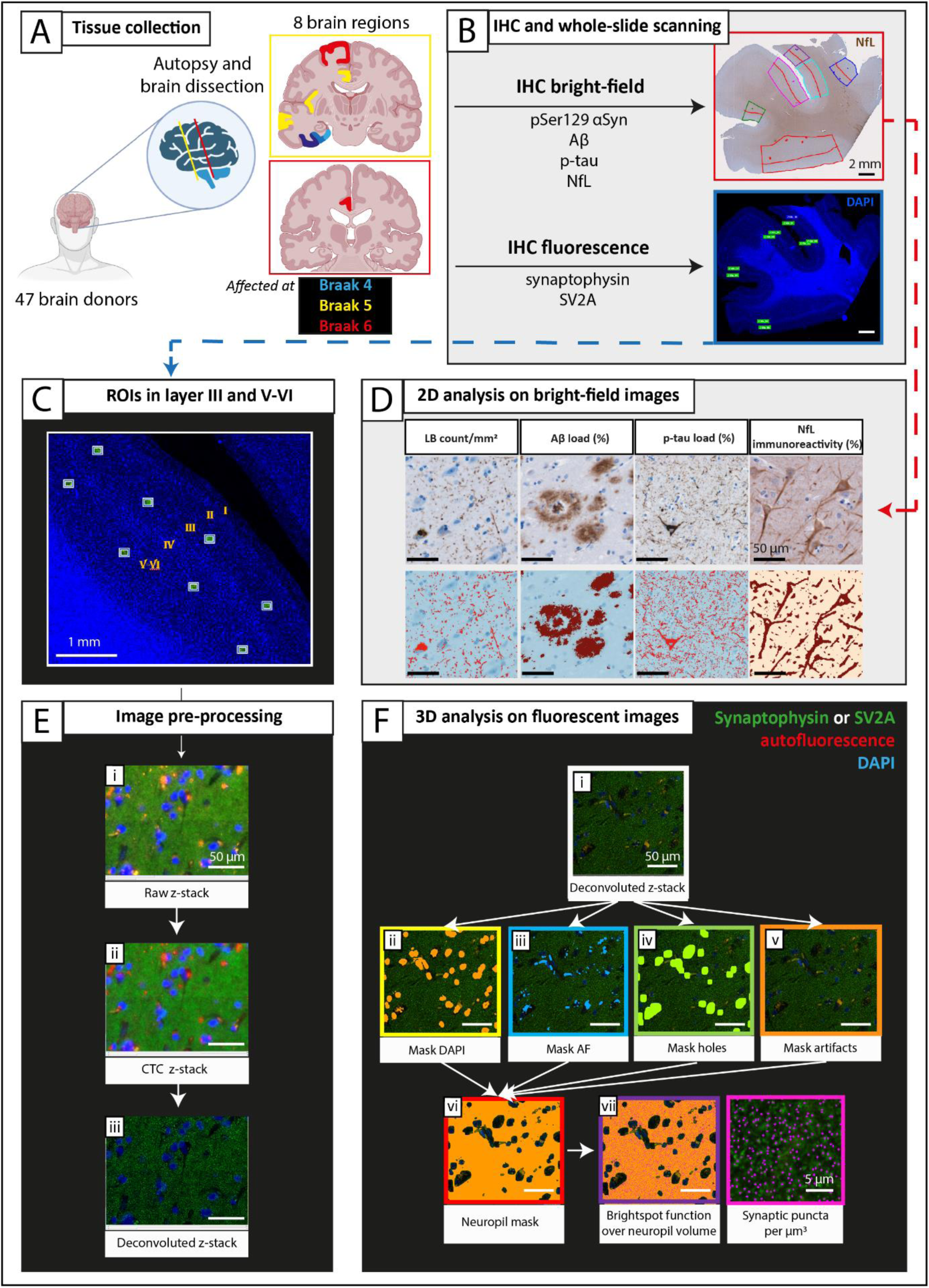
Methods workflow. **A**) After donor inclusion, autopsy was performed, and 8 cortical regions were dissected and grouped into regions affected at [41] Braak α-synuclein stage 4 (in blue), 5 (in yellow), and 6 (in red). **B)** Brain tissue was processed for immunohistochemistry using antibodies targeting pSer129 α-synuclein, p-tau, Aβ, and NfL with bright-field, and synaptophysin and SV2A with fluorescence. Whole tissue slides were digitalized, and regions of interest (ROIs) were drawn in superficial (layer I-III) and deep (layer IV-VI) layers of the cortex on bright-field scans, and in the corresponding layer III and layer V-VI on fluorescent scans (detailed ROI placing within cortical minicolumns in **C** and **Fig. S1**)). **D)** Subsequently, pixel and object classifiers in QuPath [38] were used to quantify neuropathology load and NfL immunoreactivity on bright-field scans. The ROIs drawn on fluorescent scans were **E)** pre-processed in Huygens Professional, and **F)** analysed with NIS elements to quantify synaptophysin^+^ and SV2A^+^ puncta per μm^3^ in the cortical neuropil volume. For example of artifact removal see **Fig. S2**. **Legend**: *Aβ: amyloid-β; CTC: cross-talk correction; IHC: immunohistochemistry; LB: Lewy body; NfL: neurofilament light chain; pSer129-αSyn: α-synuclein phosphorylated at Ser129; p-tau: phosphorylated tau; ROI: region of interest; SV2A: synaptic vesicle glycoprotein 2A*.

### Immunohistochemistry

6µm-thick sections from tissue blocks of the above-mentioned regions were cut and mounted on superfrost+ glass slides (Thermo Scientific, USA). All sections were stained for synaptophysin (C-terminal), SV2A (clone EPR23500-32), pSer129-αSyn (clone EP1536Y), amyloid-β (clone 4G8), p-tau (clone AT8) and neurofilament light chain (NfL, amino acid sequence 1-376) (for information on primary antibodies and dilutions see **Table S2**). Briefly, the sections were deparaffinised, immersed in buffer, and heated to 100°C in a steam cooker for 30 minutes for antigen retrieval. The sections were blocked for endogenous peroxidase using 1% hydrogen peroxide in tris buffered saline (TBS; pH 7.4), and consequently in 3% normal donkey serum in TBS (Triton 0.5%). For the synaptic stainings, Triton was not used to avoid synaptic vesicles to be washed away. Primary antibodies were added, and the sections incubated overnight at 4°C. Primary antibodies were detected using secondary antibodies labelled to Alexa 488, and the sections were counterstained with DAPI and mounted with Mowiol (Sigma-Aldrich, St. Louis, United States) plus anti-fading agent DABCO. For bright-field stainings (pSer129-αSyn, amyloid-β, p-tau, and NfL), primary antibodies were diluted in 1% normal donkey serum in TBS (Triton 0.1%) and incubated overnight at 4°C. Primary antibodies were detected using EnVision (Dako, Glostrup, Denmark), and visualized using 3.3’-Diaminobenzidine (DAB, Dako) with Imidazole (50 mg DAB, 350 mg Imidazole and 30 uL of H_2_O_2_ per 100 mL of Tris-HCl 30 mM, pH 7.6). The sections were counterstained with haematoxylin, after which were dehydrated and mounted with Entellan (Merck, Darmstadt, Germany). In between steps, TBS was used to wash the sections.

### Image analysis

#### Microscopy of synaptic staining

Synaptic imaging was performed using the Olympus VS200 (Evident). First, an overview fluorescence scan was performed using a 10× objective (NA 0.40) on the DAPI channel (**Figure 1B**). Once the overview scans were obtained, 10 regions of interest (ROIs) (200 x 160 μm) were placed within the cortex. Specifically, 5 ROIs were placed in cortical layer III, and 5 ROIs in cortical layers V-VI. Each pair of ROIs (one in cortical layer III, and one in cortical layer V-VI) was placed within a cortical minicolumn (**Fig. 1C**, for details see **Fig. S1**), corresponding to the superficial and deep pyramidal layers of the cortex, commonly affected in PD(D) and DLB [10, 15]. To ensure correspondence between ROIs for pathological/neurofilament and synaptic measurements, fluorescence ROIs were placed within corresponding sections of bright-field ROIs (**Figure 1B**, for details see **Fig. S3**). Slides were scanned overnight with 60× oil objective (NA 1.42) on 3 channels: DAPI, Alexa 488 (targeting synaptophysin or SV2A) and autofluorescence (see **Table S3** for scanning details). Images were taken as z-stacks to allow deconvolution and 3D image analysis. Specifically, 2 images were acquired above and under the focus point, with a step size of 0.28 μm in the z-direction. The scans were visually inspected for artifacts, and if needed, re-scans were immediately performed.

#### Image processing and 3D analysis of synaptic markers

Image pre-processing was performed with Huygens Professional (https://svi.nl/Huygens-Professional). First, cross-talk correction (CTC) was performed for subtraction of the autofluorescence signal (such as blood vessels, lipofuscin and artifacts, **Figure 1E, ii**), and next color deconvolution was performed (**Figure 1E, iii**). Subsequently, NIS-elements AR analysis version 5.42.00 (https://www.microscope.healthcare.nikon.com/products/software/nis-elements) was used to quantify synaptic density over volume (**Figure 1F**). Images were masked to define the neuropil volume by applying and inverting masks for 1) nuclei on the DAPI channel (**Figure 1F, ii**), 2) blood vessels on the autofluorescence channel (**Figure 1F, iii**), 3) holes in the tissue on the Alexa 488 channel (**Figure 1F, iv**), 4) and tissue artifacts such as foldings and bubbles on the Alexa 488 channel (**Figure 1F, v**; for an example of artifact removal see **Fig. S2**). Finally, all masks were inverted and combined (**Figure 1F, vi**) to define the neuropil volume in which synaptic puncta were counted. As synaptic diameter varies approximately between 0.1-1 µm [37], we incrementally inputted a diameter from 0.1 to 1 µm to identify the synaptic puncta. 0.6 µm and 0.5 µm were the diameters that recognized all synaptophysin-positive (synaptophysin+) and SV2A-positive (SV2A+) puncta, respectively. Synaptic puncta were finally counted with the ‘bright spots’ function, based on diameter and intensity (**Figure 1F, vii**). The outcome measures were number of synaptophysin^+^ and SV2A^+^ puncta/µm^3^ per ROI, yielding a total of ∼3700 data points for each synaptic marker across all cases and regions. Extreme outliers (which lie more than 3 times the interquartile range) were quality checked and excluded if necessary.

#### Quantification of NfL and pathological hallmarks

Using a whole-slide scanner (Vectra Polaris, 20× objective) images of NfL, pSer129 α-synuclein, amyloid-β and p-tau immunostained sections were taken and quantified using QuPath 0.2.3 stardist (https://qupath.readthedocs.io/en/0.2/) [38]. ROIs containing all cortical layers were delineated in straight areas of the cortex to avoid over-or underestimation of pathology in sulci and gyri, respectively [39], as described before [29, 40]. The cortex was further segmented into superficial (I-III) and deep (IV-VI) cortical layers based on the haematoxylin counterstaining. The entorhinal cortex, parahippocampal gyrus and fusiform gyrus were segmented as described before [29] (for details, see **Fig. S3**). Amyloid-β and p-tau stainings of control cases were delineated in superficial (layer I-III) and deep cortical layers (layer IV-VI) only in entorhinal, parahippocampal and fusiform gyrus. DAB immunoreactivity was quantified with in-house QuPath scripts, using pixel and object classifiers, as done previously [29]. The outcome measures for NfL was %area load, expressed in the text as %immunoreactivity. The outcome measure for pSer129-αSyn was LB density (LB count/mm^2^), while outcome measures for amyloid-β and p-tau were %area load (**Fig. 1D**).

### Statistics

Statistical analyses were performed in SPSS 28.0 (Chicago, IL). Normality was tested, and demographics between PD, PDD, DLB and control groups were compared using one-way ANOVA or Kruskal-Wallis test for continuous data, and Fisher exact test for categorical data. Since the data have a nested design (8 brain regions per donors, 10 ROIs per brain region in 2 cortical layers) multi-level linear mixed models (LMM) were used to compare synaptic densities between groups, with case ID, brain area and cortical layer as nested variables, and with age and sex as covariates. In order to associate synaptic density with regional neuropathological load, we averaged the 5 measurements in cortical layer III and V-VI to have one measure for superficial and deep cortical layers, respectively, corresponding to the data format of neuropathological load, and therefore corresponding the synaptic and neuropathological data within the same ROIs. A logarithmic transformation (log10) was applied to measures of LB density, amyloid-beta load and p-tau load after adding a constant (+1) to avoid zero to become undefined values. To assess associations across all regions between synaptic density and neuropathology load or NfL immunoreactivity, synaptic density was the dependent variable and neuropathology load or neurofilament immunoreactivity (independent variable) were the main effects, together with age and sex (i.e. covariates). The intercept was included in all analyses as random effect. Multiple pairwise (post-hoc) comparison corrections were performed using Bonferroni post-hoc test, and multiple testing comparison corrections were performed with Bonferroni method, after which p-values < 0.05 were considered significant. Group comparisons were expressed as percentage differences, as in the formula: %difference = [(absolute difference)/mean control]*100. Correlation coefficients (r) of LMM associations were calculated as in the formula: r = (estimate fixed effect * standard deviation fixed effect) / standard deviation dependent variable. Finally, we predicted synaptic density by adding all variables in the model (age, sex, neuropathological load and NfL immunoreactivity), always including the intercept as random effect.

## Results

### Cohort description

Clinical and neuropathological data of included PD, PDD, DLB and non-neurological control donors are summarized in **Table 1** (and per donor in **Table S1**). Sex, age at diagnosis, and post-mortem delay did not differ between groups. PD donors were significantly older at death than controls (mean 82 vs 72 years, *p=0.016*). Disease duration was significantly shorter for DLB compared to PD donors (mean 5 *vs* 18 years, *p<0.001*) and PDD (mean 5 *vs* 14 years, *p=0.012*). CDR scores were significantly higher in PDD (*p=0.024*) and DLB (*p=0.006*) compared to PD donors. Pathologically, DLB donors showed significantly more AD co-pathology compared to controls (ABC score: *p<0.002,* Thal phase: *p=0.010,* Braak NFT stage: *p<0.001),* but not compared to PD and PDD cases (*p>0.05*). Moreover, DLB cases had significantly more CAA type 1 compared to the other groups (DLB: 50% of cases had CAA type 1; controls: 5%, PD: 0%, and PDD: 11%, *p=0.004)*. Compared to controls, Braak NFT stage was also significantly higher in PD (*p=0.007*) and PDD (*p=0.048*). For representative images of all outcome measures see **Fig. 2**, and for quantitative regional cortical neuropathology burden and NfL immunoreactivity see **Fig. S4.**

**Figure 2.**
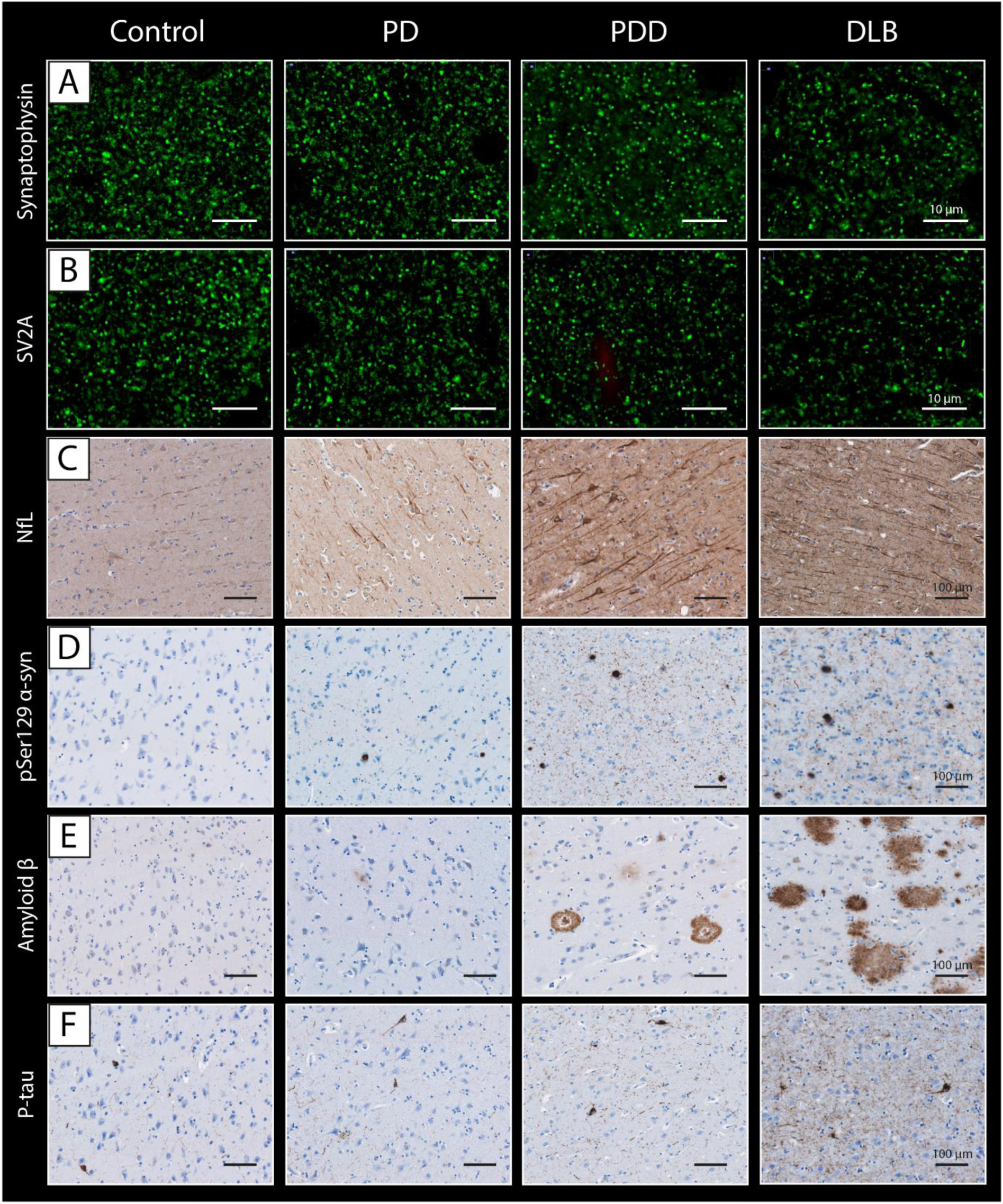
Representative images of synaptic puncta, NfL and pathological hallmarks in controls, PD, PDD, and DLB donors. Representative images of **A)** synaptophysin^+^ pre-synaptic puncta, **B)** SV2A^+^ pre-synaptic puncta, **C)** NfL, **D)** pSer129 α-synuclein, **E)** amyloid-beta and **F)** p-tau were acquired in the parahippocampal gyrus of controls, PD, PDD and DLB donors. **Legend**: *DLB: Dementia with Lewy Bodies; NfL: neurofilament light chain; PD: Parkinson’s disease; PDD: Parkinson’s disease dementia; PHG: parahippocampal gyrus*.

**Table 1.** Demographic, clinical and pathological characteristics of brain donors. Data are mean, and minimum to maximum range. Legend: CAA: cerebral amyloid angiopathy; DLB: dementia with Lewy bodies; F: females; hrs: hours; LB: Lewy body; M: males; MDI: motor-to-dementia interval; min: minutes; n.a.: not available; NFT: neurofibrillary tangles; PD: Parkinson’s disease; PDD: Parkinson’s disease dementia. \**p<0.05*, \*\**p*<0.01, \*\*\**p<0.001* when compared to controls; ^Ϯ^*p<0.05*, ^Ϯ^ ^Ϯ^*p*<0.01, ^Ϯ^ ^Ϯ^ ^Ϯ^*p<0.001* when compared to PD; ^#^*p<0.05,* ^##^*p*<0.01, ^###^*p<0.001* when compared to PDD.

|  | Control | PD | PDD | DLB |
| --- | --- | --- | --- | --- |
| **N** | 20 | 12 | 9 | 6 |
| **Sex F/M (%F)** | 11/9 (55%) | 3/9 (25%) | 5/4 (56%) | 1/5 (17%) |
| **Age at diagnosis**<br>years, mean [range] | - | 64<br>[54-76] | 63<br>[44-84] | 73<br>[62-88] |
| **Disease duration**<br>years, mean [range] | - | 18<br>[13-23] | 14<br>[8-21] | 5<sup>†††</sup> ##<br>[3-7] |
| **MDI**<br>years, mean [range] | - | - | 10<br>[2-16] | 0<sup>†††</sup><br>[0-0] |
| **CDR**<br>median (N) [range] | *n.a.* | 0.5 (N=5)<br>[0.5-0.5] | 2<sup>††</sup> (N=6)<br>[1-3] | 2<sup>†</sup> (N=5)<br>[1-3] |
| **Age at death**<br>years, mean [range] | 72<br>[57-87] | 82 *<br>[69-93] | 77<br>[62-94] | 76<br>[61-91] |
| **Post-mortem delay**<br>hrs:min, mean [range] | 8:38<br>[4:35-12:40] | 7:31<br>[3:00 -10:40] | 7:53<br>[5:10-10:40] | 7:21<br>[4:30-10:10] |
| **Pathological characteristics** |  |  |  |  |
|  | **Control** | **PD** | **PDD** | **DLB** |
| **Braak α-synuclein stage [10] N** | 20 | 12 | 9 | 6 |
| 0/1/2/3/4/5/6 | 17/2/1/0/0/0/0 | 0/0/0/0/1/1/10 | 0/0/0/0/0/0/9 | 0/0/0/0/0/0/6 |
| **ABC score [42] N** | 20 | 12 | 9 | 6 |
| A 0/1/2/3 | 4/14/2/0 | 1/8/3/0 | 0/4/3/2* | 0/1/2/3 ** |
| B 0/1/2/3 | 4/15/1/0 | 0/8/3/0 | 0/6/3/0 | 0/1/4/1 ** |
| C 0/1/2/3 | 20/0/0/0 | 11/1/0/0 | 6/1/2/0 | 1/0/4/1 *** ††† |
| **Thal phase [43] N** | 20 | 12 | 9 | 6 ** |
| 0/1/2/3/4/5 | 4/9/5/1/4/0 | 1/5/3/3/0/0 | 0/3/1/3/2/0 | 0/1/0/2/1/2 |
| **Braak NFT stage [44] N** | 20 | 12 * | 9 * | 6 *** |
| 0/1/2/3/4/5/6 | 4/11/4/1/0/0/0 | 0/3/5/3/1/0/0 | 0/2/4/1/2/0/0 | 0/0/1/1/3/0/1 |
| **CAA type [45] N** | 20 | 12 | 9 | 6<sup>†</sup> |
| No CAA/type 1/ type 2 | 16/1/3 | 11/0/1 | 6/1/2 | 2/3**/1 |

Data are mean, and minimum to maximum range. Legend: CAA: cerebral amyloid angiopathy; DLB: dementia with Lewy bodies; F: females; hrs: hours; LB: Lewy body; M: males; MDI: motor-to-dementia interval; min: minutes; n.a.: not available; NFT: neurofibrillary tangles; PD: Parkinson's disease; PDD: Parkinson's disease dementia. \* $p < 0.05$ , \*\* $p < 0.01$ , \*\*\* $p < 0.001$ when compared to controls; $^{\dagger}p < 0.05$ , $^{\dagger\dagger}p < 0.01$ , $^{\dagger\dagger\dagger}p < 0.001$ when compared to PD; $^{\#}p < 0.05$ , $^{\#\#}p < 0.01$ , $^{\#\#\#}p < 0.001$ when compared to PDD.

### Regional synaptophysin and SV2A loss in PD, PDD and DLB

Synaptophysin density was decreased by 4% in PD (*p=0.048*), 7% in PDD (*p<0.001*), and 8% in DLB (*p<0.001*) compared to controls across all regions **(Fig. 3A).** When looking at regions affected at different Braak α-synuclein stages, synaptophysin density was significantly lower in regions affected at Braak 4 (i.e. entorhinal cortex, parahippocampal gyrus and fusiform gyrus combined) in DLB compared to controls (–9%, *p=0.011*). Additionally, synaptophysin density was significantly lower in regions affected at Braak 5 (i.e. middle temporal gyrus, anterior cingulate cortex and anterior insula combined) in PDD versus controls (–11%, *p<0.001*) and versus PD (–7%, *p=0.022*) **(Fig. 3C, E).** Synaptophysin density did not differ between groups in regions affected at Braak 6 (i.e. superior frontal gyrus and posterior cingulate gyrus combined). In summary, synaptophysin density was decreased in all groups compared to controls, and more specifically in regions affected at Braak 4 in DLB, and in regions affected at Braak 5 in PDD.

**Figure 3.**
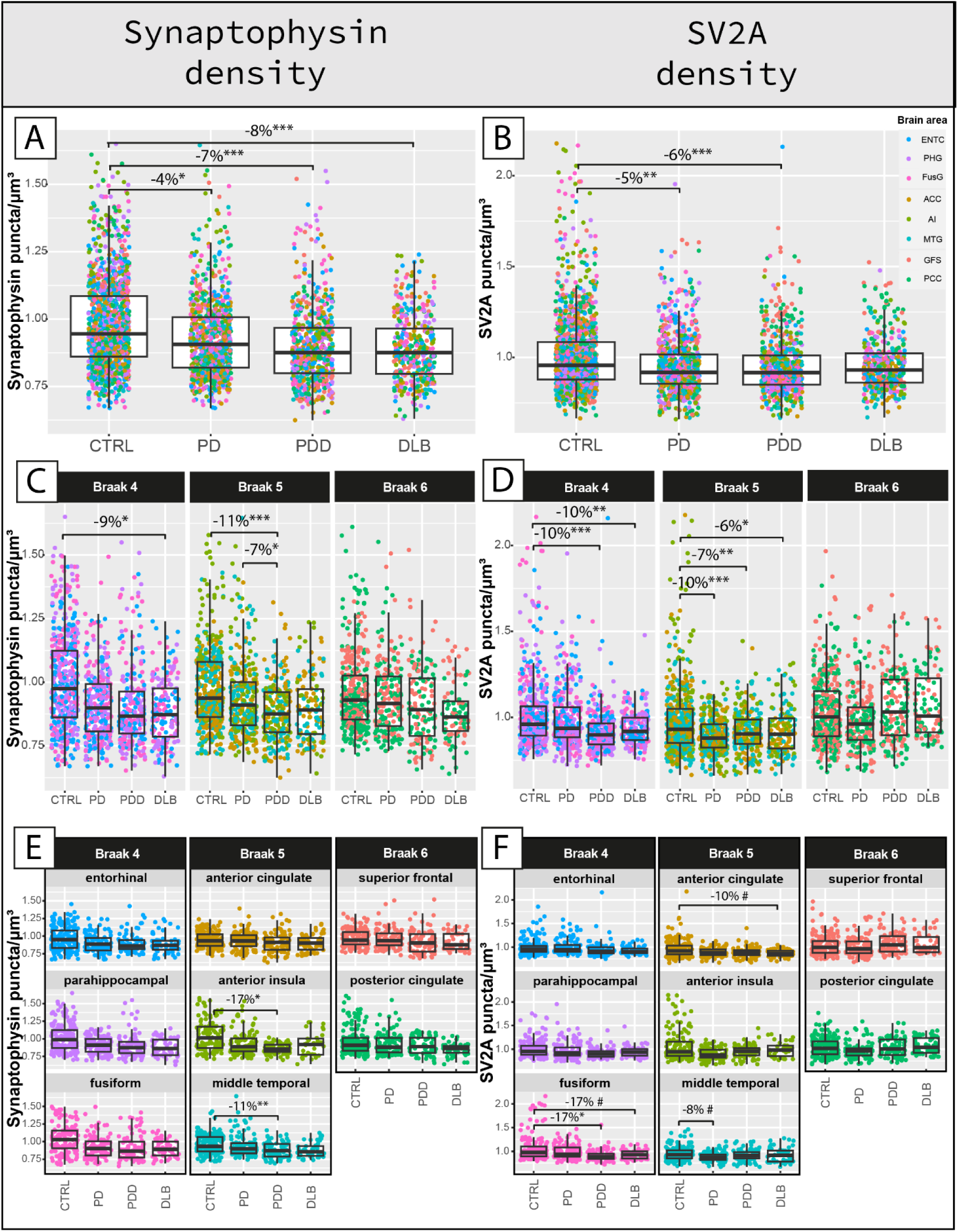
Synaptophysin and SV2A density are reduced in PD, PDD and DLB in a region-specific manner compared to controls. **A,B**) The *first row* shows the group differences in synaptophysin and SV2A density across all cases, **C,D)** the *second row* shows the same group differences within regions grouped per brain areas affected at Braak 4, 5 and 6, and **E, F)** the *last row* shows the same group differences within brain areas. Every data point represents one measurement color-coded by brain area. Linear mixed model p-values are adjusted for multiple pairwise (post-hoc) comparisons between groups and multiple testing comparisons for regions affected at Braak stage 4, 5, and 6 in **C)** and **D)**, and for 8 regions in **E)** and **F)**. ^#^p<0.100, *p<0.05, **p<0.01, ***p<0.001 after Bonferroni post-hoc test. **Legend**: *ACC: anterior cingulate gyrus; AI: anterior insula; CTRL: control; DLB: Dementia with Lewy Bodies; ENTC: entorhinal cortex; FusG: fusiform gyrus; GFS: superior frontal gyrus; MTG: middle temporal gyrus; PCC: posterior cingulate gyrus; PD: Parkinson’s disease; PDD: Parkinson’s disease dementia; PHG: parahippocampal gyrus.* ^#^*p<0.10,* \**p<0.05*, \*\**p*<0.01, \*\*\**p<0.001*.

SV2A density was decreased by 5% in PD (*p=0.006*), and 6% in PDD (*p<0.001*), and did not differ in DLB (*p=0.105*) compared to controls across all regions **(Fig. 3B).** When looking at regions affected at different Braak α-synuclein stages, SV2A density was significantly lower in regions affected at Braak 4 in DLB (–10%, *p=0.001*) and PDD (–10%, *p<0.001*) compared to controls, and in regions affected at Braak 5 in PD (–10%, *p<0.001*), PDD (–7%, *p=0.003*) and DLB (–6%, *p=0.047*) compared to controls **(Fig. 3D, F).** SV2A density did not differ between groups in regions affected at Braak 6. In summary, SV2A density was decreased in all groups compared to controls, specifically in regions affected at Braak 4 in DLB and PDD (especially the fusiform gyrus), and in regions affected at Braak 5 in all groups.

Synaptic density decreased with age and was higher in females (**supplementary results** and **Fig. S5**), and was slightly higher in layer III compared to later VI (**supplementary results** and **Fig. S6**). Layer III and VI seemed to be similarly affected in PD, PDD, DLB (**Fig. S7**). Within-groups, synaptophysin density was similar across brain regions, while SV2A density was significantly higher in regions affected at Braak stage 6 in PD, PDD and DLB (**supplementary results** and **Fig. S8**). As such, regional synaptophysin and SV2A density only weakly correlated (**supplementary results** and **Fig. S9**).

### Cortical synaptic loss is associated with NfL immunoreactivity

We previously showed that NfL accumulates in cortical brain regions in PDD and DLB compared to controls, and that increased NfL immunoreactivity can be considered a proxy of axonal fragmentation and degeneration [29]. Here, we investigated whether cortical synaptic density correlated with regional NfL immunoreactivity.

Both synaptophysin (r = –0.16, *p<0.001*) and SV2A density (r = –0.08, *p=0.033*) negatively correlated with NfL immunoreactivity across all cases. Regionally (**Fig. 4A**), synaptophysin density showed a strong negative correlation with NfL immunoreactivity specifically in the regions affected at Braak 6 in DLB cases (r = –0.68, R^2^=46%, *p=0.048*), and not in any region of any other groups (*p*>0.05). Regarding SV2A density (**Fig. 4B**), a significant negative correlation with NfL immunoreactivity was found in the regions affected at Braak 4 of PD cases (r = –0.35, R^2^=12%, *p=0.030*), and at Braak 6 of DLB cases (r = –0.83, R^2^=69%, *p<0.001*).

**Figure 4.**
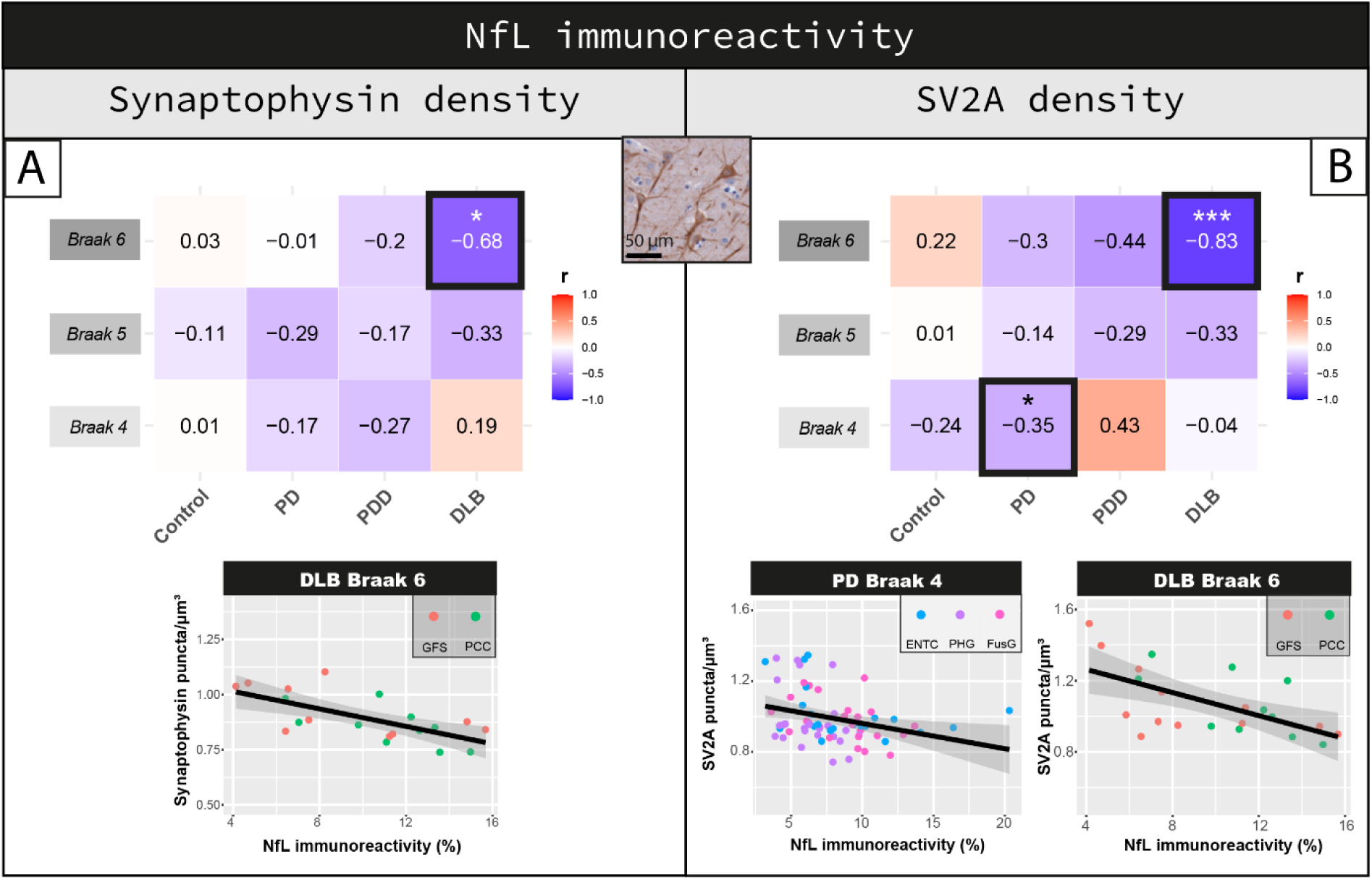
Inverse correlation between synaptophysin or SV2A and NfL density. Correlation of NfL immunoreactivity with **(A)** synaptophysin and **(B)** SV2A density are shown per group in regions affected at Braak 4, 5 and 6. Heat-maps are color-coded for correlation coefficient (r), being blues representative of negative correlations and reds of positive correlations. For each correlation, the correlation coefficient is reported, together with its p-value. For the correlations highlighted with a square we are also reporting the scatterplots, color-coded for brain area (every data point represents one averaged measurement per layer III and V-VI). Linear mixed model p-values are adjusted for multiple testing (12 correlations per synaptic marker) *p<0.05, **p<0.01, ***p<0.001 after Bonferroni post-hoc test. **Legend**: *CTRL: control; DLB: Dementia with Lewy Bodies; ENTC: entorhinal cortex; FusG: fusiform gyrus; GFS: superior frontal gyrus; NfL: neurofilament light chain; PCC: posterior cingulate gyrus; PD: Parkinson’s disease; PDD: Parkinson’s disease dementia; PHG: parahippocampal gyrus*.

Taken together, SV2A density associated with increased NfL immunoreactivity in PD and DLB. Particularly, regions affected at Braak 6 of DLB cases showed the strongest correlation where NfL immunoreactivity accounted for up to 70% of the variability in SV2A density.

### Synaptic loss is more severe in cortical brain regions with higher LB density

Synaptophysin density (PD+PDD+DLB: r = –0.12, R^2^=1%, *p=0.006*) and SV2A (PD+PDD+DLB: r = –0.18, R^2^=3%, *p<0.001*), negatively correlated with LB density across all regions in the disease groups. Regionally (**Fig. 5A**), synaptophysin density negatively correlated with LB density in regions affected at Braak 4 in PD (r = –0.46, R^2^=21%, *p=0.011*) and DLB groups (r = –0.48, R^2^=23%, *p=0.005*), and in regions affected at Braak 6 in PDD (r = –0.55, R^2^=30%, *p=0.039*). Synaptophysin density did not correlate with LB density in any other region (*p*>0.05). Moreover, SV2A density correlated negatively with LB density in regions affected at Braak 4 in PD (r = –0.39, R^2^=15%, *p=0.022*), but not in other regions or groups (**Fig. 5B**).

**Figure 5.**
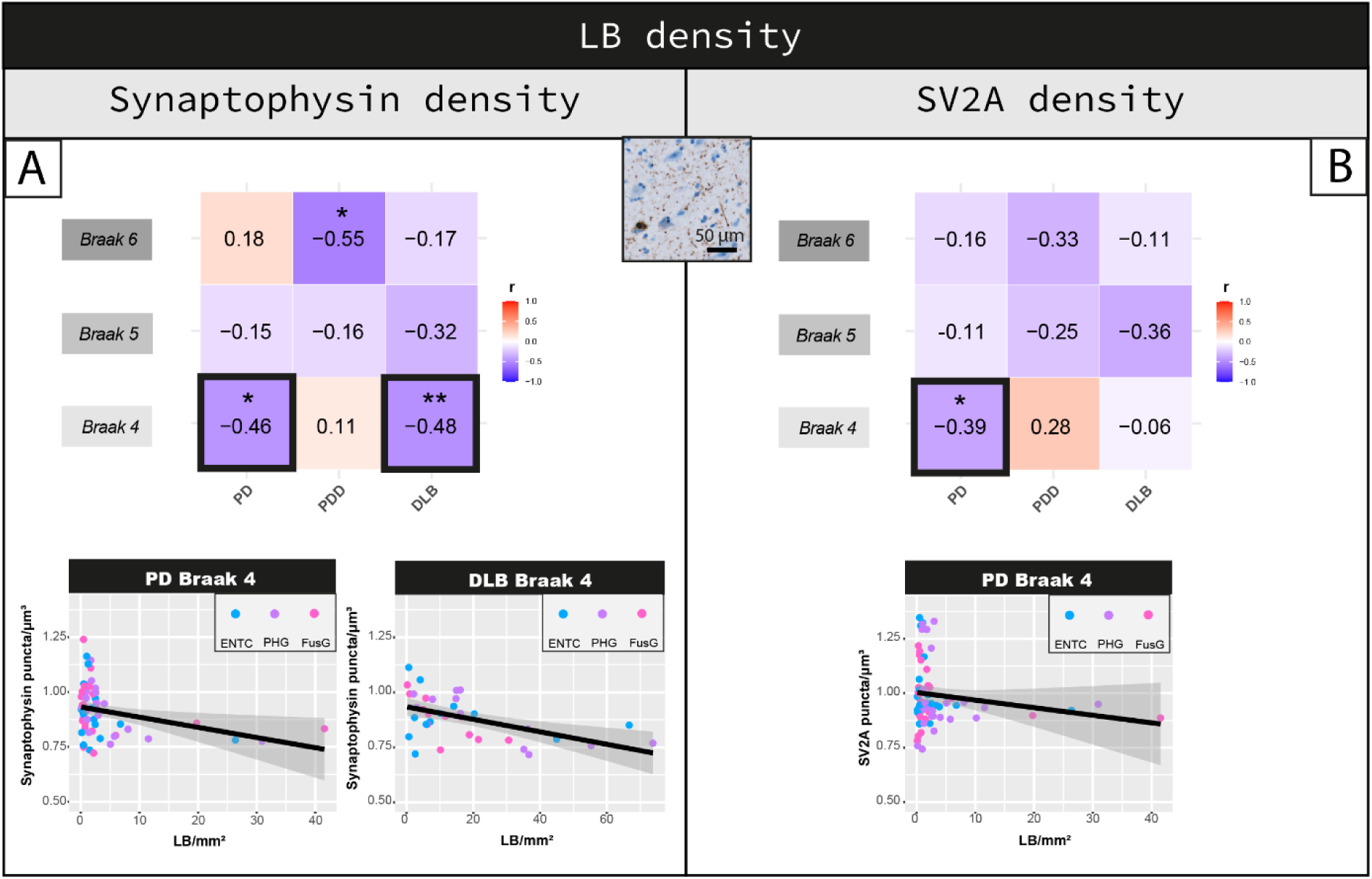
Cortical synaptophysin and SV2A density shows an inverse correlation with LB density in regions affected at Braak stage 4 in PD and DLB. Correlation between LB density and **(A)** synaptophysin and **(B)** SV2A density are shown per groups per regions affected at Braak 4, 5 and 6. Heat-maps are color-coded for correlation coefficient (r), being blues representative of negative correlations and reds of positive correlations. For each correlation, the correlation coefficient is reported, together with its p-value. Linear mixed model p-values are adjusted for multiple testing (9 correlations per synaptic marker). *p<0.05, **p<0.01, ***p<0.001 after Bonferroni post-hoc test. For the correlations highlighted with a square we are also reporting the scatterplots, color-coded for brain area (every data point represents one averaged measurement per layer III and V-VI). **Legend**: *CTRL: control; DLB: Dementia with Lewy Bodies; ENTC: entorhinal cortex; FusG: fusiform gyrus; GFS: superior frontal gyrus; LB: Lewy body; PCC: posterior cingulate gyrus; PD: Parkinson’s disease; PDD: Parkinson’s disease dementia; PHG: parahippocampal gyrus*.

Taken together, increased LB density associated with synaptophysin and SV2A loss mostly in regions affected at Braak 4, which are the regions showing the highest LB density **(Fig. S4)**.

### Higher SV2A density is linked to higher amyloid-β load

SV2A density, but not synaptophysin (*p=0.453*), positively correlated with amyloid-β load across all regions of all cases (r = 0.14, R^2^=2%, *p=0.001*). Regionally, synaptophysin density negatively correlated with amyloid-β load in regions affected at Braak 4 of DLB cases (r = – 0.52, R^2^=27%, *p=0.023*) (**Fig. 6A**). As a strong negative association was found in these regions with LB pathology (see previous paragraph), we corrected for LB density in the analysis. Indeed, decreased synaptophysin density in regions affected at Braak 4 tended to be driven by LB density rather than amyloid-β load (amyloid-β: *p=0.305*; LB density: *p=0.055*). SV2A density positively correlated with amyloid-β load in regions affected at Braak 6 in DLB (r = 0.76, R^2^=58%, *p<0.001*), but not in other groups (*p*>0.05) (**Fig. 6B**).

**Figure 6.**
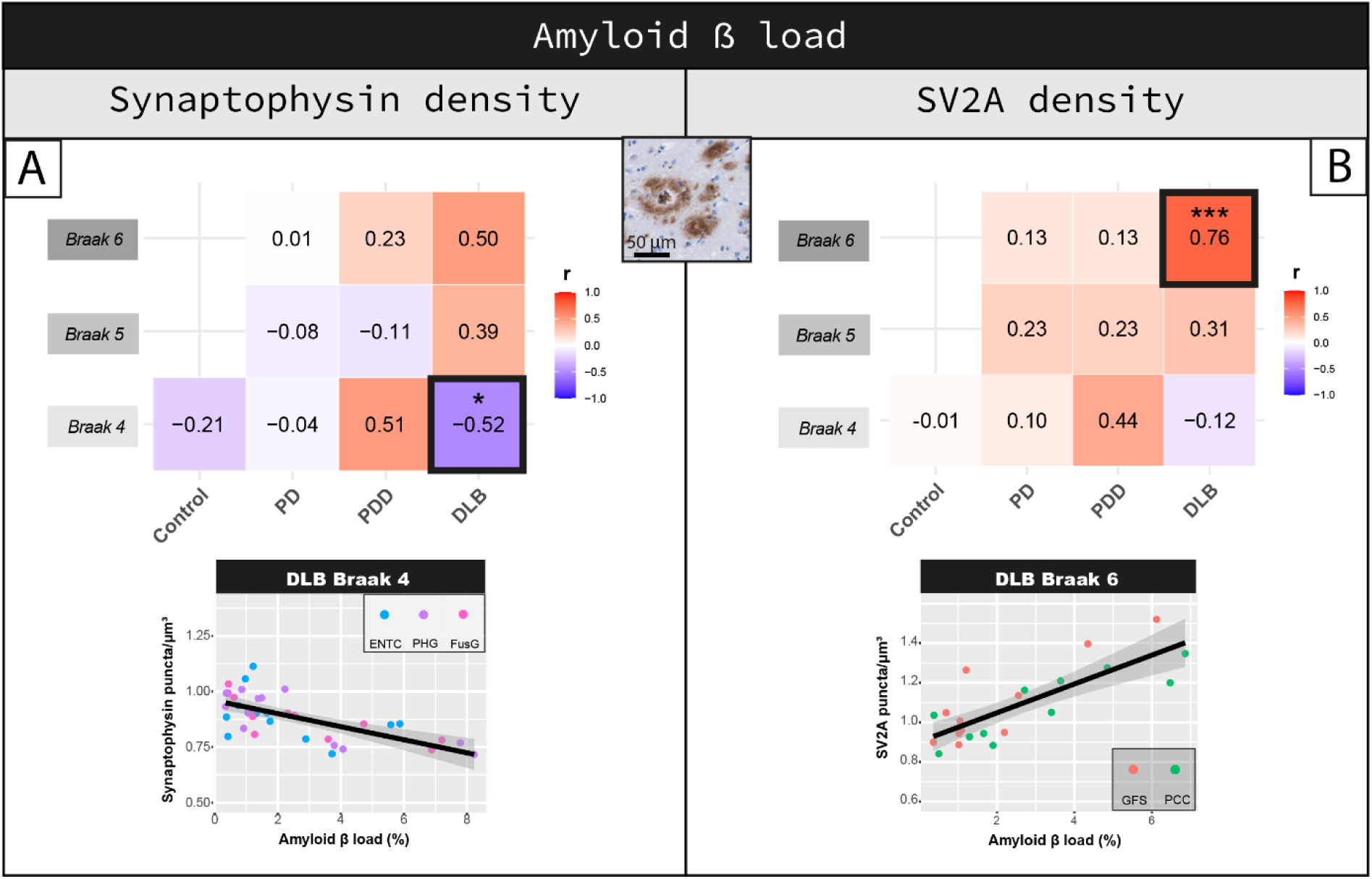
Cortical SV2A density shows a positive correlation with amyloid-β load in DLB. Correlation between amyloid-β load and **(A)** synaptophysin and **(B)** SV2A density are shown per group in regions affected at Braak 4, 5 and 6. Heat-maps are color-coded for correlation coefficient (r), being blues representative of negative correlations and reds of positive correlations. For each correlation, the correlation coefficient is reported, together with its p-value. Linear mixed model p-values are adjusted for multiple testing (10 correlations per synaptic marker). ^#^p<0.100, *p<0.05, **p<0.01, ***p<0.001 after Bonferroni post-hoc test. For the correlations highlighted with a square we are also reporting the scatterplots, color-coded for brain area (every data point represents one averaged measurement per layer III and V-VI). **Legend**: *CTRL: control; DLB: Dementia with Lewy Bodies; ENTC: entorhinal cortex; FusG: fusiform gyrus; GFS: superior frontal gyrus; PCC: posterior cingulate gyrus; PD: Parkinson’s disease; PDD: Parkinson’s disease dementia; PHG: parahippocampal gyrus*.

Taken together, higher SV2A density in DLB was associated with higher amyloid-β load in regions affected at Braak 6, where amyloid-β load accounted for almost 60% of the variability in SV2A density. No effects for synaptophysin were found after correcting for LB pathology.

### P-tau load has differential effects on synaptophysin and SV2A density

Synaptophysin density, and not SV2A (*p=0.598*), negatively correlated with p-tau load across all regions of all cases (r = –0.10, R^2^=1%, *p=0.022*). Regionally, synaptophysin density did not correlate with p-tau load in any region of any group (*p>0.05*) (**Fig. 7A**). SV2A density positively correlated with p-tau load in regions affected at Braak 4 in PDD (r = 0.85, R^2^=72%, *p<0.001*), but not in other groups (>0.05) (**Fig. 7B**). As a similar association was found in this region for amyloid-beta load (see **Fig. 6**), we assessed if this association was driven by amyloid-beta. Increased SV2A density in regions affected at Braak 4 in PDD cases was driven by an increase in amyloid-β load rather than p-tau load (amyloid-β: r = 1.00, *p<0.001;* p-tau: r= –0.74, *p=0.044)*.

**Figure 7.**
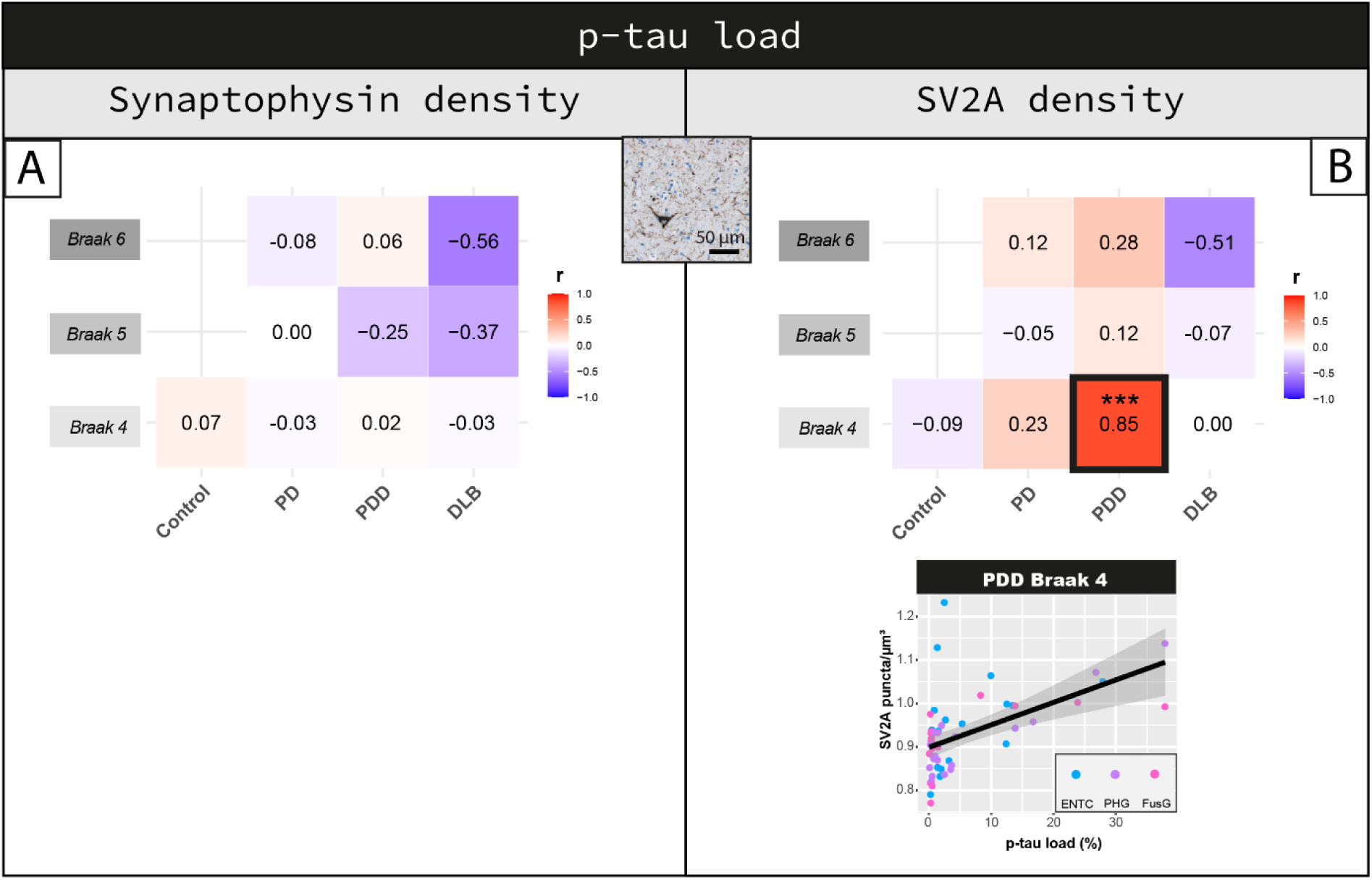
Cortical synaptophysin and SV2A density differentially correlate with p-tau load. Correlation between p-tau load and **(A)** synaptophysin and **(B)** SV2A density are shown per group in regions affected at Braak 4, 5 and 6. Heat-maps are color-coded for correlation coefficient (r), towards blue represents a negative correlations and towards red a positive correlations. For each correlation, the correlation coefficient is reported, together with its p-value. For the correlations highlighted with a square we are also reporting the scatterplots, color-coded for brain area (every data point represents one averaged measurement per layer III and V-VI). Linear mixed model p-values are adjusted for multiple testing (10 correlations per synaptic marker). *p<0.05, **p<0.01, ***p<0.001 after Bonferroni post-hoc test. **Legend**: *CTRL: control; DLB: Dementia with Lewy Bodies; ENTC: entorhinal cortex; FusG: fusiform gyrus; GFS: superior frontal gyrus; PCC: posterior cingulate gyrus; PD: Parkinson’s disease; PDD: Parkinson’s disease dementia; PHG: parahippocampal gyrus*.

Taken together, p-tau load seemed to have differential effects on synaptophysin and SV2A density. P-tau weakly correlated with synaptophysin density across all regions, while it positively associated with SV2A density in regions affected at Braak 4 in PDD, but this effect was mostly driven by amyloid-β load.

### Synaptic degeneration is linked to axonal damage, higher LB density and older age

We investigated the combined contribution of age, sex, NfL immunoreactivity, LB density, amyloid-β and p-tau load on synaptophysin and SV2A density by including all these markers in the same model (for models in regions affected at Braak stage 4, 5 and 6 see **Table S4**).

For synaptophysin, the model explained 56% of the variance (R^2^) when including all cases. The strongest negative associations were observed between synaptophysin density and LB density (β = –0.074, *p<0.001)*, NfL immunoreactivity (β = –0.008, *p=0.001)*, and age (β = –0.002, *p=0.017)*. When looking at the model including only the diseased groups (PD+PDD+DLB) (R^2^=54%), LB density (β = –0.053, *p=0.019)*, and NfL immunoreactivity (β = –0.008, *p<0.001)* were the negative predictors of synaptophysin density.

For SV2A, an explained variance of 57% was observed across all cases. The strongest negative associations were seen between SV2A density and LB density (β = –0.103, *p<0.001)*, NfL immunoreactivity (β = –0.007, *p=0.002)*, and age (β = –0.002, *p=0.021)*, and a positive association for amyloid-β load (β = 0.170, *p<0.001)*. When looking at the model including only the diseased groups (PD+PDD+DLB) (R^2^=61%), the results did not change: the negative predictors were again LB density (β = –0.130, *p<0.001*), NfL immunoreactivity (β = –0.006, *p=0.003),* and age (β = –0.004, *p<0.001),* and the only positive predictor amyloid-β load (β = 0.210, *p<0.001)*.

Taken together, both synaptophysin and SV2A densities associated with increased LB density, NfL immunoreactivity, and age. Amyloid-β load was a specific positive predictor for SV2A density. Both models explained around between 56-61% of the variance in synaptic density.

### Cortical synaptophysin loss is more severe in cases with lower cognitive scores

Lastly, we investigated whether cortical synaptophysin and SV2A density correlated with clinical measures, including disease duration, motor-to-dementia onset and CDR scores.

Longer disease duration significantly correlated with cortical synaptophysin loss in DLB cases (r = –0.27, *p=0.009*) and with cortical SV2A loss in PDD cases (r = –0.19, *p=0.015*). A longer motor-to-dementia onset correlated with cortical SV2A loss in PDD cases (r = –0.32, *p<0.001*), but not with cortical synaptophysin loss (*p>0.05*). Higher CDR scores significantly associated with cortical synaptophysin loss (r = –0.21, *p<0.001*), but not with SV2A loss (*p=0.179*).

Taken together, we found that longer disease duration, longer motor-to-dementia onset and higher CDR scores correlated with synaptic loss.

## Discussion

Using a high throughput immunofluorescence approach, we mapped and quantified synaptic loss in eight cortical brain regions commonly affected in PD(D) and DLB, and assessed whether regional differences in the pattern of synaptic loss are linked to axonal degeneration and neuropathological burden. We found that synaptic loss was most severe in temporal, cingulate and insular cortices affected at Braak stage 5 in PD, and additionally in parahippocampal regions affected at Braak stage 4 in PDD and DLB, and that synaptic loss associated with NfL immunoreactivity and LB density. Lastly, we found that cortical synaptic degeneration associated with longer disease duration and worse cognitive scores.

Both synaptophysin and SV2A density were decreased in PD, PDD and DLB compared to non-demented controls. The effect size of synaptic loss we found matches well with the literature in both post-mortem and PET studies [15, 24]. Similar to our results of a 4-8% decrease in cortical synaptic density, a post-mortem study found a global reduction of cortical synaptophysin density of 5% in PD and 8% in PDD compared to age-matched controls [15]. PET studies using a SV2A tracer also reported 3-10% decrease in binding in cortical regions of PD patients [23, 25], which was more pronounced in PDD/DLB [24]. Regionally, we found that cortical regions affected at Braak α-synuclein stage 4 and 5 were more severely affected by synaptic degeneration, while cortical regions affected at Braak α-synuclein stage 6 showed synaptic density comparable to controls. These results suggest that synaptic degeneration occurs in regions affected by α-synuclein, and since Braak α-synuclein stage also describes an increase in α-synuclein load with disease progression, it is probably dependent on the amount of α-synuclein pathology [10]. Furthermore, we found that the pattern of synaptic degeneration differed between PD and PDD/DLB cases: in PD temporal, cingulate and insular cortices affected at Braak α-synuclein stage 5 showed more pronounced synaptic degeneration compared to controls, while PDD and DLB cases additionally showed synaptic degeneration in parahippocampal regions affected at Braak α-synuclein stage 4. These results suggest that the pattern of regional synaptic neurodegeneration differs in PD and PDD/DLB, and that parahippocampal regions show more synaptic degeneration in demented cases.

Synaptophysin and SV2A densities were both decreased in PD, PDD and DLB, but they only correlated weakly between each other. Both synaptophysin and SV2A sub-cellular localization is in synaptic vesicles at the pre-synaptic compartment [46, 47]. Our study shows that they were both more abundant in layer III than V-VI of the cortex, as expected [48], and their loss weakly associated with age, as shown before [15, 18]. However, we found a different regional distribution of the two markers: while SV2A density was higher in cortical regions affected at Braak α-synuclein stage 6 in PD(D) and DLB, synaptophysin density did not show regional density differences, which goes in line with the literature [15]. Therefore, our results suggest that even if synaptophysin and SV2A both target synaptic vesicles in the pre-synaptic compartment, they might have a differential expression or involvement in disease.

Our results indicate that NfL immunoreactivity partially contributed to regional and global synaptic loss. We previously showed that NfL accumulates in parahippocampal regions in PDD/DLB, and that increased NfL immunoreactivity can be considered a proxy of axonal damage [29]. Increased NfL levels in biofluids are commonly used as a proxy of axonal degeneration in neurodegenerative disorders [30], but evidence of how NfL levels relate to synaptic markers is still limited. The only study investigating the correlation between regional SV2A PET binding and global CSF NfL levels did this in an AD cohort, and reported a negative correlation between NfL levels and SV2A binding in several cortical regions [49]. In line with these results, we show a coupling between regional synaptic degeneration and NfL accumulation in the brain of DLB donors, especially in regions affected at the end stage of disease, indicating that here synaptic degeneration might occur together with axonal degeneration.

We found that synaptic degeneration associated with higher LB density in parahippocampal regions affected at Braak stage 4. In physiological conditions, α-synuclein resides in the pre-synaptic compartment [12], where it might start aggregating during disease [1, 13, 14]. Previous research investigating the role of α-synuclein in cell and animal models showed that α-synuclein overexpression led to inhibition of neurotransmitter release, by reducing the size of synaptic vesicle recycling pool [50], finally inducing synaptic failure and loss. In addition, a post-mortem study reported a negative correlation between LB density and synaptophysin levels in the occipital cortex of DLB donors [51]. In line with these results, we show that synaptic loss is more severe in cortical regions with the highest LB density, supporting the hypothesis that severe α-synuclein accumulation and aggregation contributes to synaptic degeneration.

We also addressed whether AD co-pathology contributes to cortical synaptic loss in PD, PDD, and DLB by investigating the correlations between synaptic measures, amyloid-β and p-tau load in cortical brain regions. Cortical p-tau load associations with synaptic density showed conflicting results, and did not contribute to synaptic loss in the model testing the combined contribution of all markers. We found that higher cortical SV2A density associated with higher amyloid-β load in demented donors. These findings place themselves in a complex framework based on previous literature, as there is no clear and consistent association of amyloid-β with synaptic density. In AD mouse models, both a loss [52, 53] and a clustering [53] of synaptic puncta were described at the margins of amyloid-β plaques. In our study, in order to measure synaptic puncta only within the neuropil, we excluded DAPI positive structures, which also include compact plaques [54] **(Fig. S10)**. Therefore, our findings might relate to two phenomena: i) a rearrangement of SV2A^+^ pre-synaptic terminals near amyloid-β plaques [53], or ii) SV2A^+^ synapses being pushed by amyloid-β plaques in a more restricted neuropil volume. This last hypothesis is supported by a PET study in mild cognitive impairment and AD [55], which reported a strong positive correlation between hippocampal amyloid-β deposition and increased SV2A density. Taken together, our results suggest that severe amyloid-β pathology might lead to increased SV2A density, and as such PET studies using SV2A tracer should take into account AD co-pathology in patient selection and interpretation of their results.

Lastly, we showed that synaptophysin loss weakly associated with longer disease duration and worse cognition scores. Several other studies, ranging from proteomics, immunohistochemistry, CSF and PET, report correlations between synaptic degeneration, disease progression [25] and cognitive decline [16–18, 56, 57]. In line with these results, we found that synaptic loss in the cortex of PD, PDD and DLB donors partially associated with disease progression and worse cognition. In addition, we showed synaptic loss in demented donors in cortical regions near the hippocampus, which are regions known to play a central role in cognition [58]. As such, our work suggests that loss of synapses might be a correlate of cognitive decline in PD(D) and DLB, and that regional differences in synaptic loss might partially underlie differences in cognitive status of these patients.

There are some limitations in our study. We only focused on 8 cortical regions, while several other regions, such as the substantia nigra, locus coeruleus, striatum, occipital and primary motor cortices have also shown to be affected in PD(D) and DLB [10, 23, 24]. In addition, we measured synaptic density only in pyramidal layers III and V-VI, as these layers have been shown to have the strongest loss of synaptic puncta in PD [15], therefore our results might not be representative of the whole cortical lamina. Lastly, even if we included a total of 47 brain donors, we still had small sample sizes per group (e.g. n=6 for DLB) and did not include any incidental Lewy body cases (cases with LB pathology and no PD clinical symptoms) to investigate the earliest prodromal disease stages. Therefore, future research should investigate synaptic density and its relationship to neuropathological burden in larger and more diverse cohorts.

## Conclusions

Synaptic neurodegeneration occurs in temporal, cingulate and insular cortices in PD, as well as in parahippocampal regions in PDD and DLB. In addition, synaptic loss was linked to axonal damage and severe α-synuclein burden. These results together with the association of synaptic loss with disease progression and cognitive impairment, indicate that regional synaptic loss may underlie clinical heterogeneity in PD(D) and DLB. Our findings might provide useful information for the interpretation of synaptic biomarkers *in vivo*.

## Declarations

### Ethical Approval and Consent to participate

All donors signed an informed consent for brain donation and the use of material and clinical information for research purposes. The procedures for brain tissue collection of NBB and NABCA have been approved by the Medical Ethical Committee of Amsterdam UMC, Vrije Universiteit Amsterdam.

### Consent for publication

Not applicable.

### Availability of supporting data

Supporting data include supplementary tables, supplementary figures, and supplementary results.

### Authors’ contributions

I.F contributed to experimental concept and design, data collection, statistical analysis, interpretation of the data, and drafting of the manuscript; M.M.A.B., R.T.G.M.M.N., and E.P. contributed to data collection; M.P. and E.T. supported the technical development of the synaptic imaging and analysis; A.J.M.R contributed to the neuropathological characterization of the cohort; W.D.J.B. and L.E.J contributed to the experimental concept and design, interpretation of the data, revised the manuscript and obtained the funding. All authors read and approved the final manuscript.

## Supporting information

Supplementary results

## Abbreviations

Aβ: amyloid-beta
ACC: anterior cingulate gyrus
AD: Alzheimer’s disease
AI: anterior insula
CSF: cerebrospinal fluid
DLB: dementia with Lewy bodies
ENTC: entorhinal cortex
FDR: false discovery rate
FusG: fusiform gyrus
GFS: superior frontal gyrus
IHC: immunohistochemistry
LB: Lewy body
MTG: middle temporal gyrus
NfL: neurofilament light chain
NFT: neurofibrillary tangles
PCC: posterior cingulate gyrus
PD: Parkinson’s disease
PDD: Parkinson’s disease dementia
PHG: parahippocampal gyrus
p-tau: phosphorylated-tau
pSer129-αSyn: phosphorylated Ser129 α-synuclein

## Acknowledgements

We would like to thank all brain donors and their next of kin for brain donation. We would also like to thank the autopsy teams of the Netherlands Brain Bank (NBB) and Normal Aging Brain collection Amsterdam (NABCA).

