## Supplementary figures for "Regional differences in synaptic degeneration are linked to alpha-synuclein burden and axonal damage in Parkinson’s disease and Dementia with Lewy bodies"

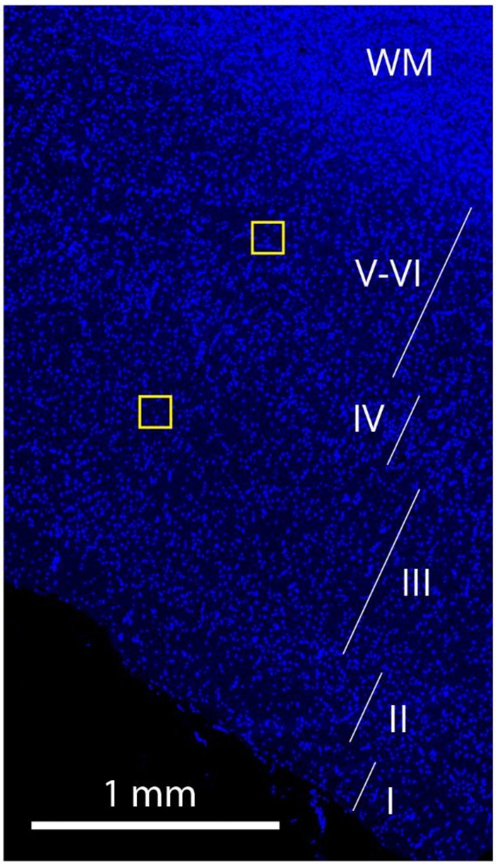


**Fig. S1 Cortical layers and placement of ROIs within minicolumns**. Cortical layers (I-VI) were clearly distinguishable in the DAPI overview, up to the start of the white matter (WM). ROIs were placed in layer III and layer V-VI within the same mini-column.

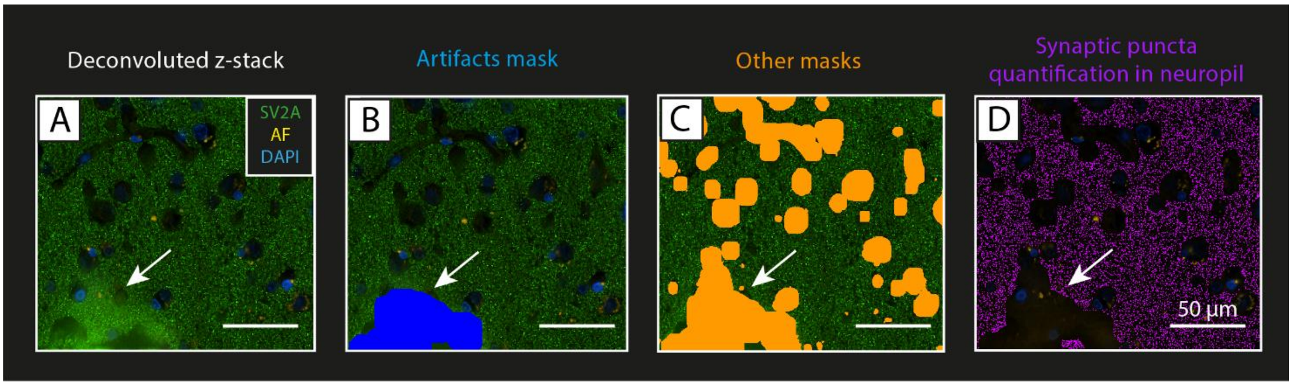


**Fig. S2 Example of artifact removal by masking. (A)** shows a deconvoluted image of the SV2A staining, with SV2A+ puncta in green, DAPI (nuclei) in blue, and autofluorescence (AF) signal in yellow. This image came with an artifact (arrow), which caused an increased signal intensity in the bottom left of the image. **(B)** In the analysis pipeline, we thresholded and masked the synaptic green signal to identify areas with high intensity signal, therefore creating an ‘artifacts mask’ (in blue, arrow). **(C)** The artifacts mask was added to the other masks (in orange, arrow) based on DAPI, autofluorescence and holes in the tissue (see **Fig. 1** for details), so that it would be excluded from the neuropil mask. **(D)** Finally, the bright-spot function was run on the neuropil mask (inverted compared to **(C)**), and SV2A+ puncta (in purple) were counted over the neuropil volume excluding also the artifact (arrow indicates artifact excluded). **Legend**: *AF: autofluorescence.*

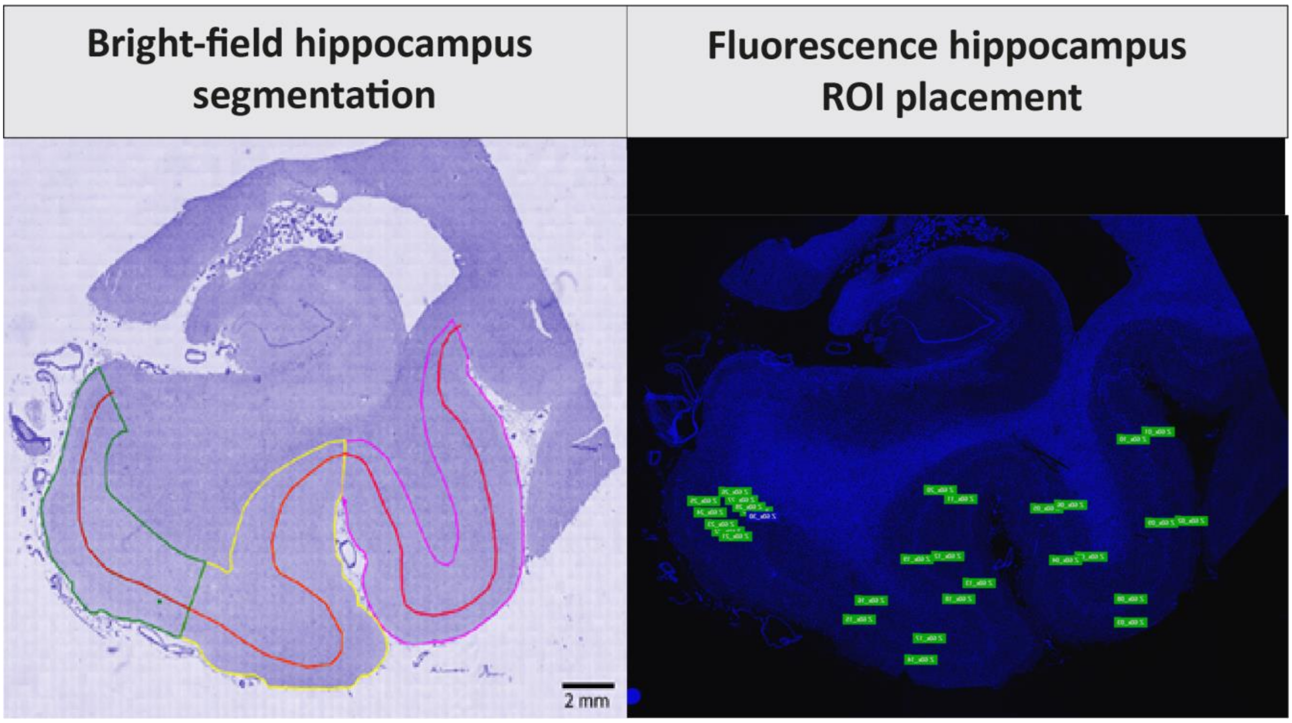


**Fig. S3 Bright-field and fluorescence hippocampal segmentation of entorhinal cortex, parahippocampal and fusiform gyrus**. Hippocampal sections were segmented according to the method described by Adler et al. [1] (on the left), where the entorhinal cortex (green), parahippocampal gyrus (yellow) and fusiform gyrus (pink) were delineated based on hematoxylin signal, and separated at the interface between layer III and VI (red line), to create superficial (I-III) and deep cortical layers ROIs (IV-VI). Briefly, in bright-field stainings (left) the entorhinal cortex was delineated from the end of the parasubiculum until layer IV started being visible [2]; the parahippocampal gyrus started from this point and ended at the collateral sulcus; the fusiform gyrus started at this point and ended at the inferior temporal sulcus. Note that the entorhinal cortex was subdivided into layers I-III (superficial) and lamina dissecans plus layers V-VI (deep) [35]. The same was applied on fluorescence stainings (right), where the ROIs (in green) were placed on the DAPI overview in layer III and V-VI within the limits of the ROIs based on the corresponding bright-field staining segmentation.

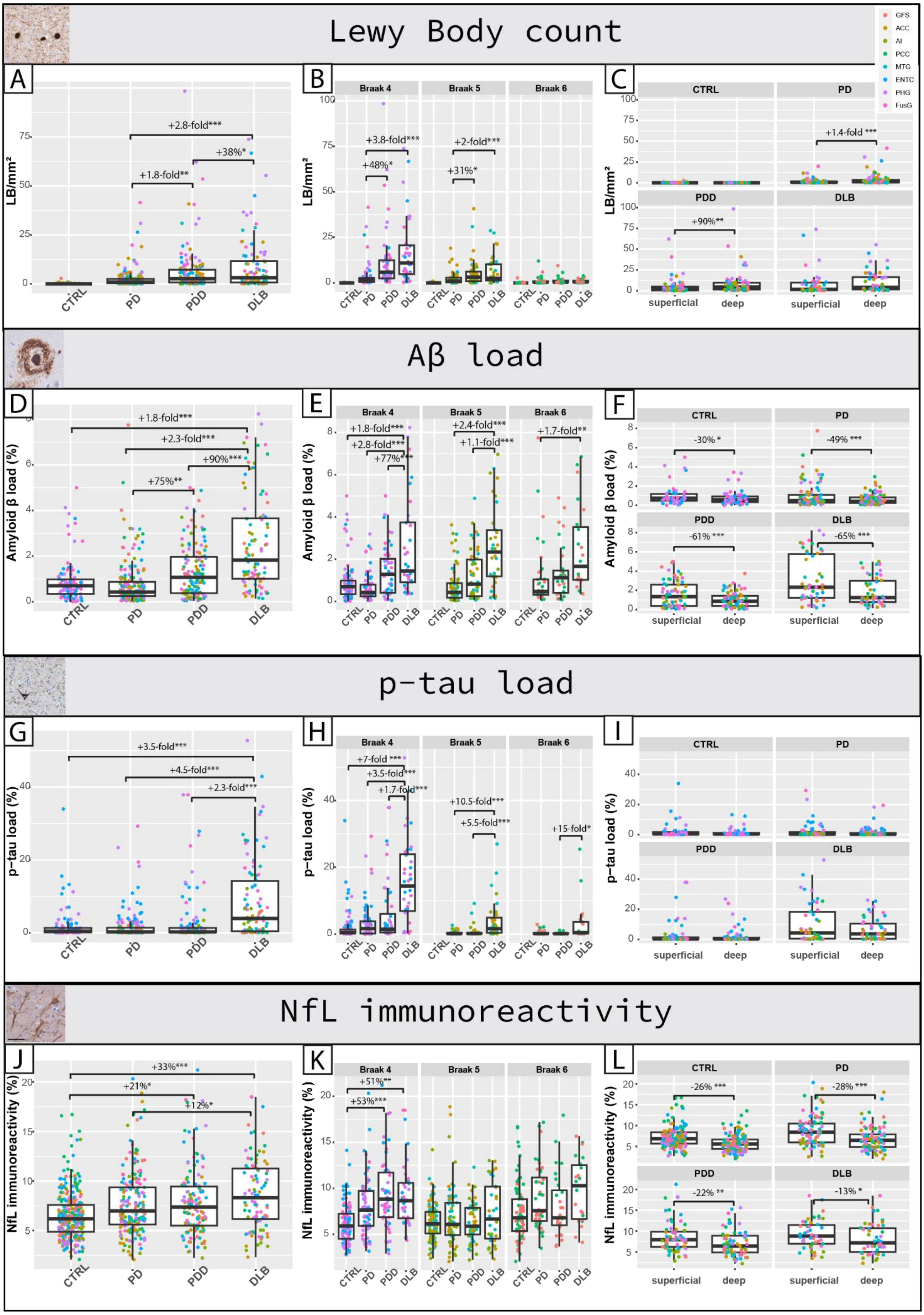

Lewy Body count
GFS
ACC
AI
PCC
MTG
ENTC
PHG
FusG
A
LB/mm²
+2.8-fold***
+38%*
+1.8-fold**
CTRL
PD
PDD
DLB
B
Braak 4
Braak 5
Braak 6
+3.8-fold***
+48%*
+2-fold***
+31%*
C
CTRL
PD
PDD
DLB
+1.4-fold ***
+90%**
superficial
deep
Aβ load
D
Amyloid β load (%)
+1.8-fold***
+2.3-fold***
+90%***
+75%**
E
+1.8-fold***
+2.8-fold***
+77%***
+2.4-fold***
+1.1-fold***
+1.7-fold**
F
-30% *
-49% ***
-61% ***
-65% ***
p-tau load
G
p-tau load (%)
+3.5-fold***
+4.5-fold***
+2.3-fold***
H
+7-fold ***
+3.5-fold***
+1.7-fold***
+10.5-fold***
+5.5-fold***
+15-fold*
I
NfL immunoreactivity
J
NfL immunoreactivity (%)
+33%***
+21%*
+12%*
K
+51%**
+53%***
L
-26% ***
-28% ***
-22% **
-13% *

**Fig. S4 Regional neuropathology load and NfL immunoreactivity. A, B** and **C** show the pSer129-αSyn-positive LB density (LB/mm$^2$), **D, E** and **F** the amyloid-β (Aβ) load (%area), **G, H** and **I** the p-tau load (%area), and **J, K** and **L** the NfL immunoreactivity (%area) for controls, PD, PDD, and DLB groups. The *left column* shows the overall pathology load across all cortical regions examined; every data point represents a single measurement (all regions, superficial and deep layers), and is color-coded based on brain area indicated in the top right of the graph. The *middle column* shows the regional pathology load in regions affected at Braak 4, 5, and 6 across groups. The *right column* shows the pathology load in superficial (I-III) and deep cortical layers (IV-VI) in each group for each outcome measure. **F** and **I** show amyloid-β and p-tau load in superficial (I-III) and deep cortical layers (IV-VI) of entorhinal, parahippocampal and fusiform cortex only. Group differences are expressed as percentage differences when under 1-fold difference, and as fold differences when over 1-fold difference. **Legend**: **p<0.05, **p<0.01, ***p<0.001; Aβ: amyloid beta; ACC: anterior cingulate cortex; AI: anterior insula; DLB: Dementia with Lewy Bodies; ENTC: entorhinal cortex; FusG: fusiform gyrus; LB: Lewy body; MTG: middle temporal gyrus; NfL: neurofilament light chain; PCC: posterior cingulate gyrus; PHG: parahippocampal gyrus; PD: Parkinson's disease; PDD: Parkinson's disease dementia; p-tau: phosphorylated tau; GFS: superior frontal gyrus.*

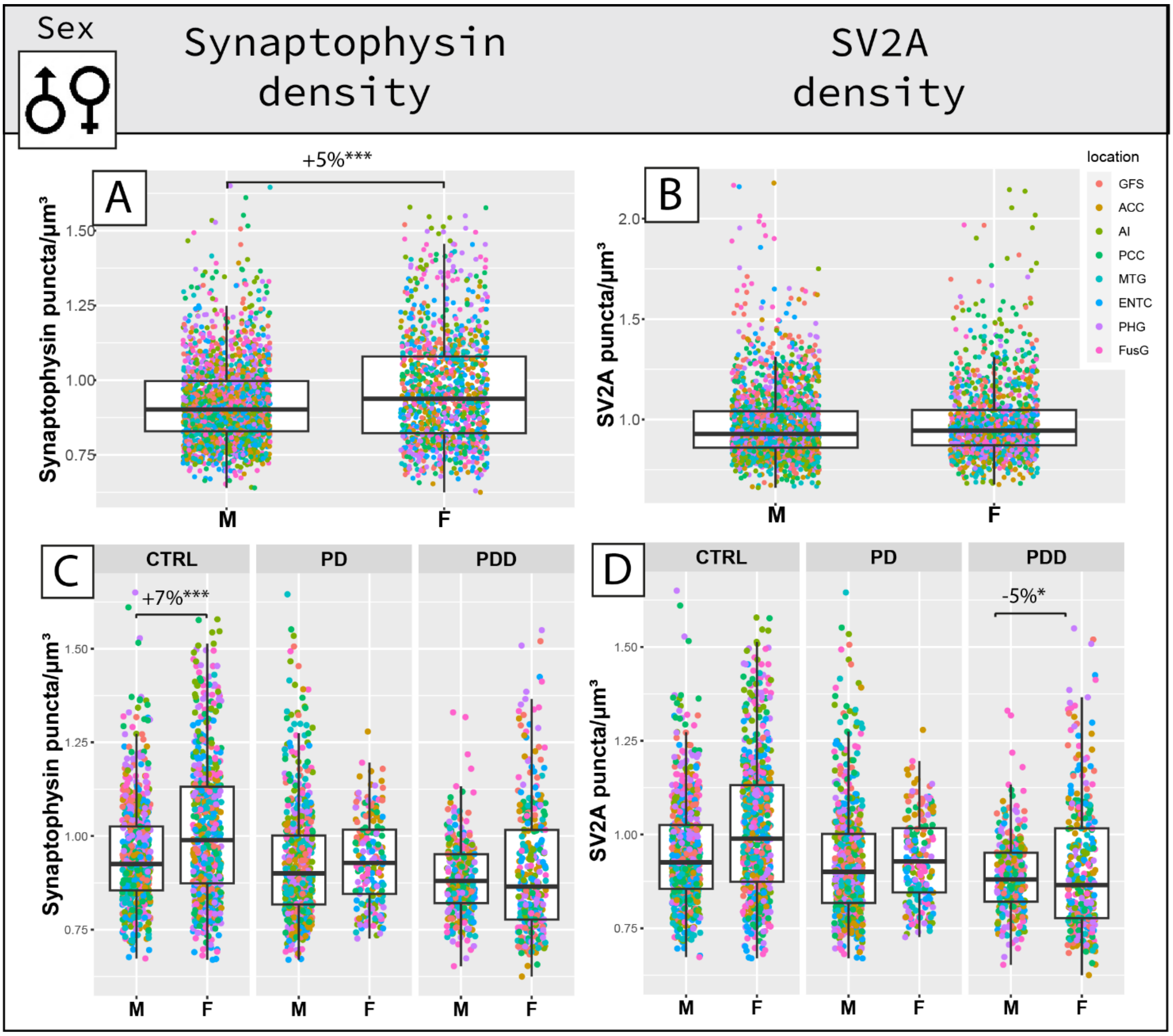


**Fig. S5 Sex differences in synaptic density.** The *top row* shows overall sex differences in synaptophysin and SV2A density, and the *bottom row* the same within groups. **(A)** Synaptophysin density was significantly higher in females (F) than males (M), **(C)** specifically by 7% in controls. **(B)** SV2A density overall did not differ in males and females, **(D)** while in the PDD group it was lower by 5% in males compared to females. **Legend**: *ACC: anterior cingulate gyrus; AI: anterior insula; CTRL: control; DLB: Dementia with Lewy Bodies; ENTC: entorhinal cortex; F: female; FusG: fusiform gyrus; GFS: superior frontal gyrus; M: male; MTG: middle temporal gyrus; PCC: posterior cingulate gyrus; PD: Parkinson's disease; PDD: Parkinson's disease dementia; PHG: parahippocampal gyrus. *p<0.05, **p<0.01, ***p<0.001.*

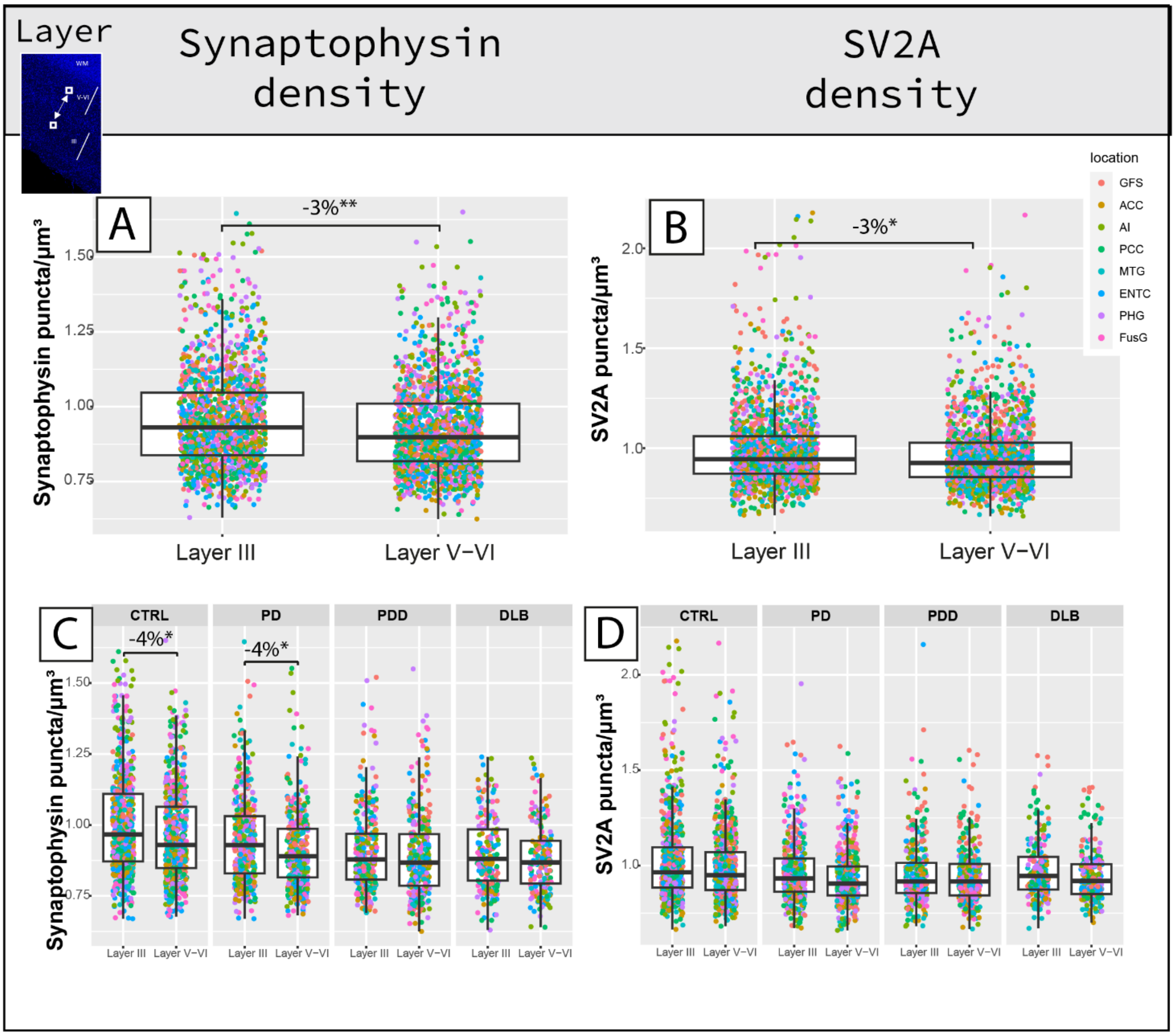


**Fig. S6 Layer III and V-VI differences in synaptic density.** The *top row* shows overall layer differences in synaptophysin and SV2A density, and the *bottom row* the same within groups. **(A)** Synaptophysin density was a significant 3% lower in layer V-VI than layer III, **(C)** specifically in controls and PD. **(B)** Similarly, SV2A density was a significant 3% lower in layer V-VI than layer III, **(D)** which was not specific to any group. **Legend**: *ACC: anterior cingulate gyrus; AI: anterior insula; CTRL: control; DLB: Dementia with Lewy Bodies; ENTC: entorhinal cortex; F: female; FusG: fusiform gyrus; GFS: superior frontal gyrus; M: male; MTG: middle temporal gyrus; PCC: posterior cingulate gyrus; PD: Parkinson's disease; PDD: Parkinson's disease dementia; PHG: parahippocampal gyrus. *p<0.05, **p<0.01, ***p<0.001.*

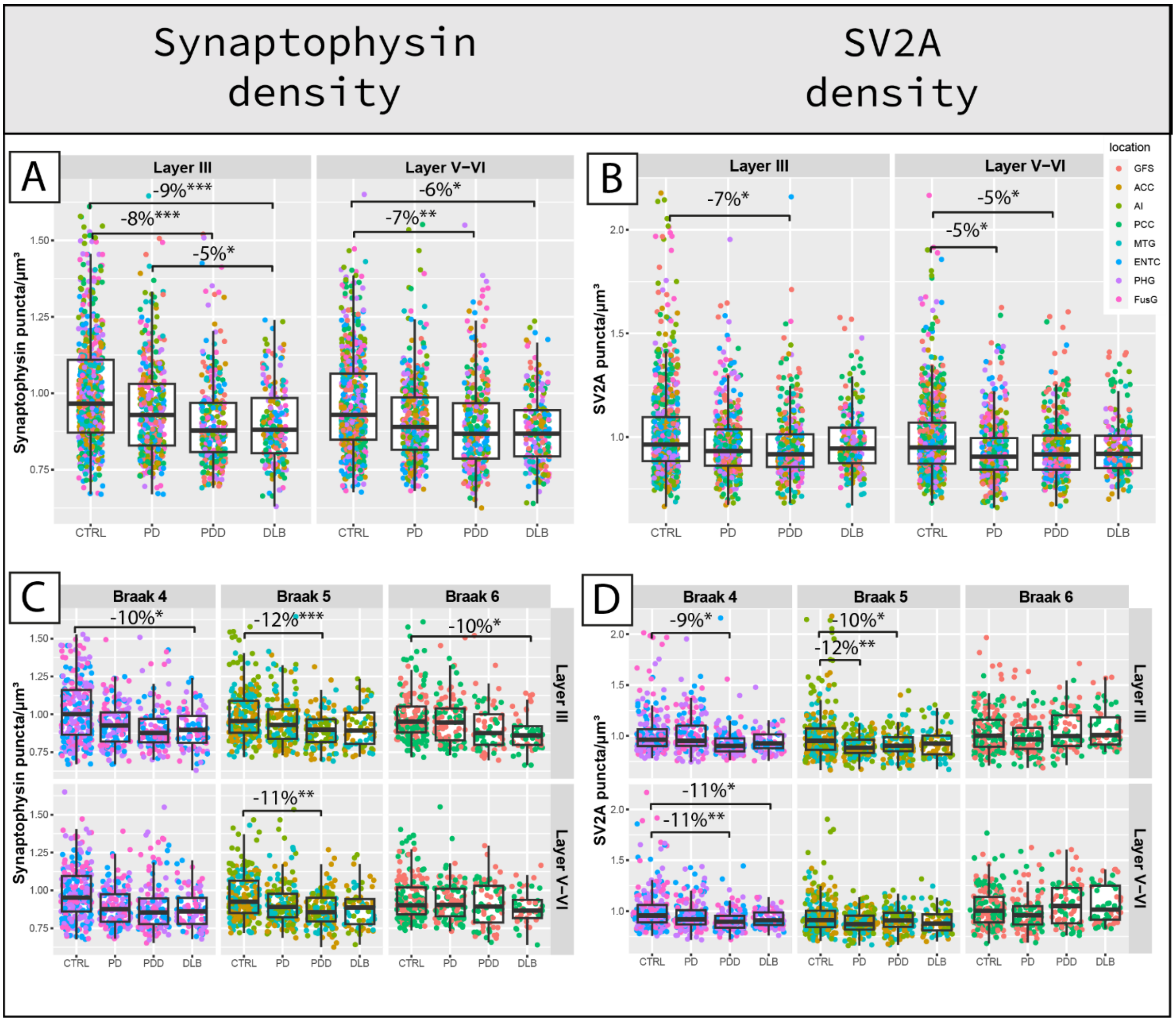


**Fig. S7 Synaptophysin and SV2A density overall and regional differences in layer III and V-VI between groups.** The *top row* shows group differences across all regions in **(A)** synaptophysin and **(B)** SV2A density within cortical layer III and V-VI. The *bottom row* shows the regional group differences in **(C)** synaptophysin and **(D)** SV2A density within cortical layer III and V-VI. Every data point represents one measurement and it is color-coded by brain area (legend top right). **Legend**: *ACC: anterior cingulate gyrus; AI: anterior insula; CTRL: control; DLB: Dementia with Lewy Bodies; ENTC: entorhinal cortex; FusG: fusiform gyrus; GFS: superior frontal gyrus; MTG: middle temporal gyrus; PCC: posterior cingulate gyrus; PD: Parkinson's disease; PDD: Parkinson's disease dementia; PHG: parahippocampal gyrus. *p<0.05, **p<0.01, ***p<0.001.*

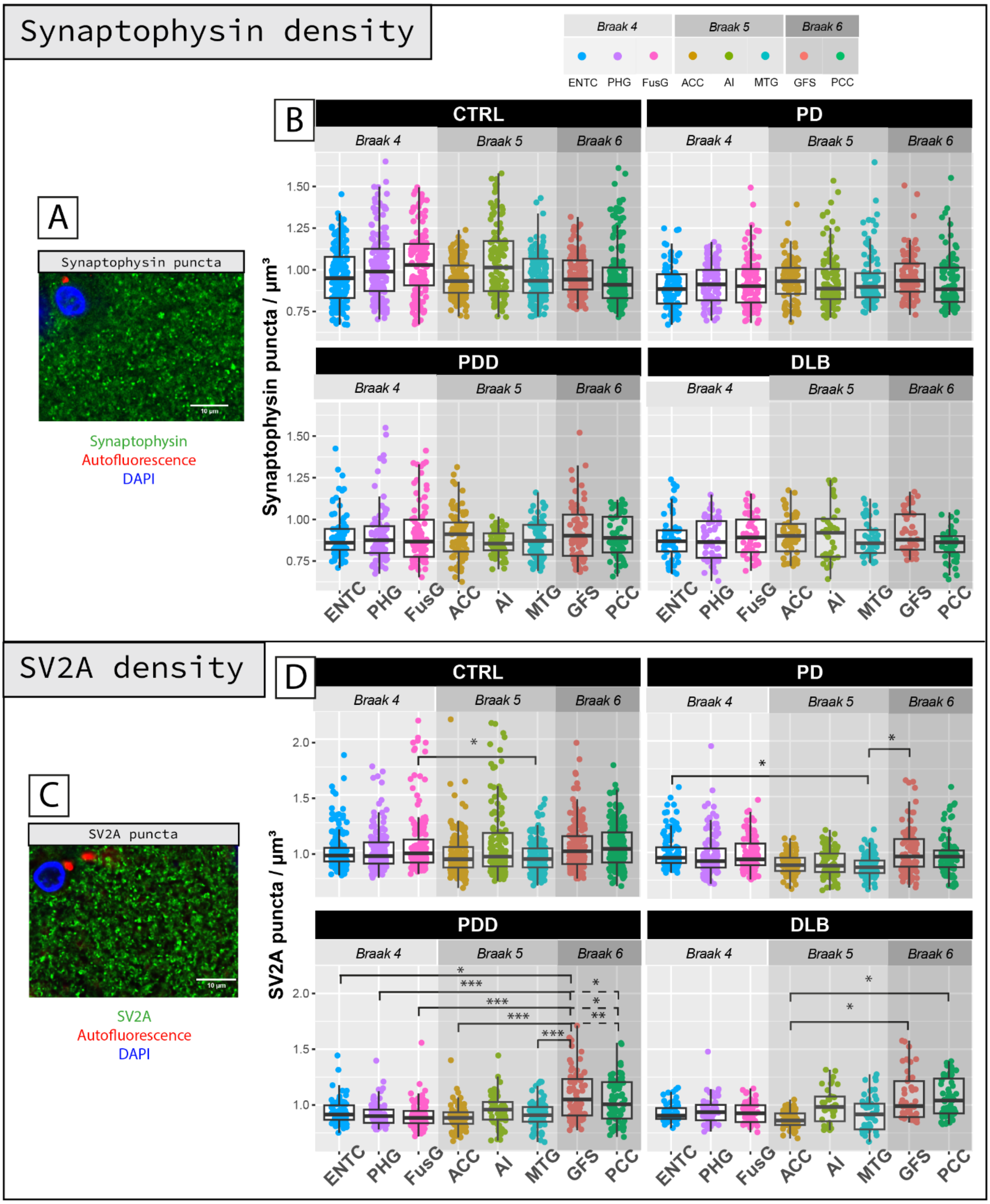


**Fig. S8 Distribution of cortical synaptophysin and SV2A density across brain regions within PD, PDD, DLB and control groups. A,C)** Synaptophysin and SV2A staining pattern in the cortex (in green; scale bar 10 µm). **B)** Synaptophysin density was similar across brain areas in the different groups. **D)** SV2A density differed across brain areas in all group: regions affected at Braak stage 6 seemed to be spared during disease compared to regions affected at earlier stages. Every data point represents one measurement color-coded by brain area. Linear mixed model p-values are adjusted for multiple testing (8 brain areas). #p<0.10, *p<0.05, **p<0.01, ***p<0.001. **Legend:** *ACC: anterior cingulate gyrus; AI: anterior insula; CTRL: control; DLB:*

*Dementia with Lewy Bodies; ENTC: entorhinal cortex; F: female; FusG: fusiform gyrus; GFS: superior frontal gyrus; M: male; MTG: middle temporal gyrus; PCC: posterior cingulate gyrus; PD: Parkinson's disease; PDD: Parkinson's disease dementia; PHG: parahippocampal gyrus.*

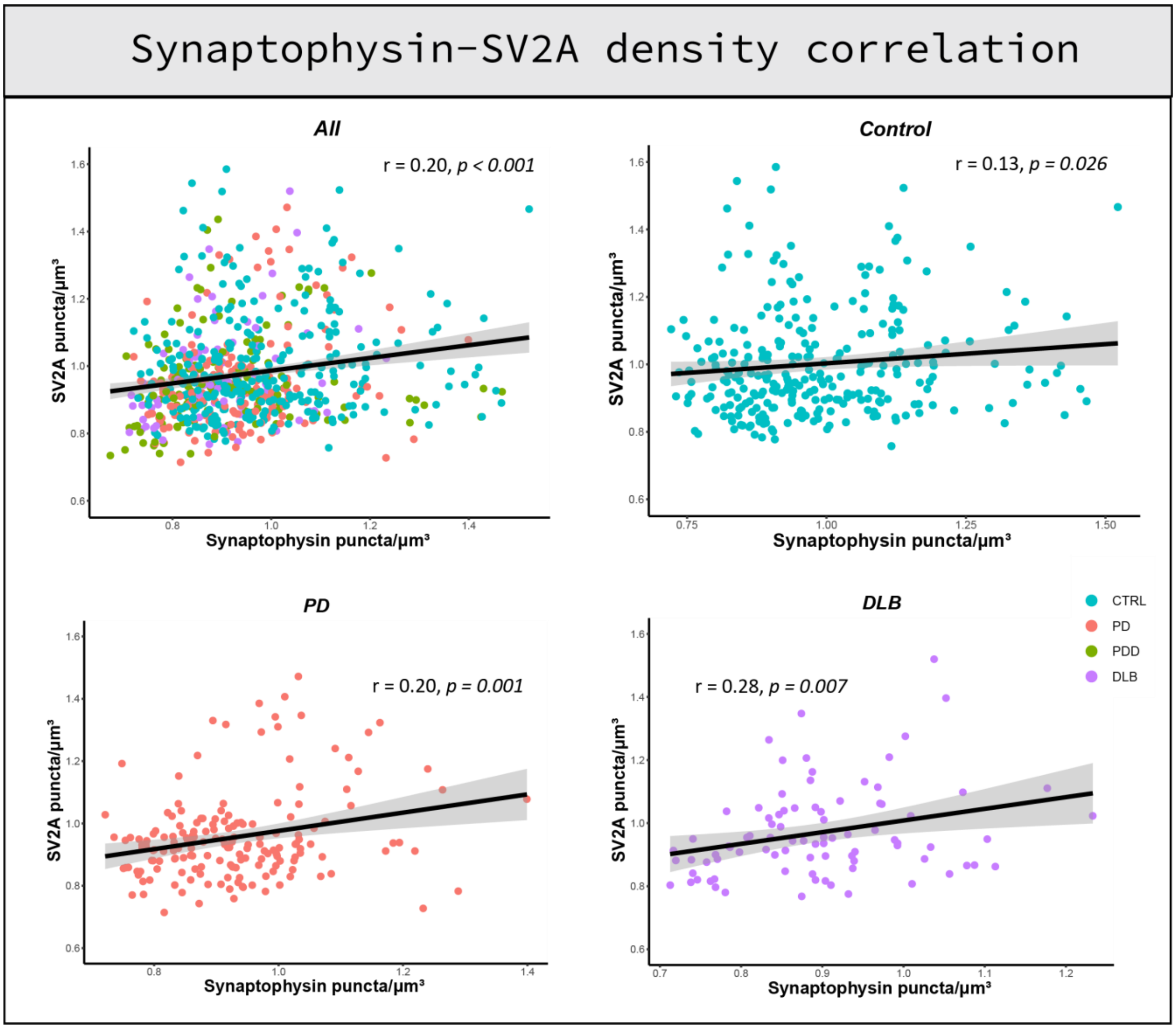


**Fig. S9 Correlation between synaptophysin and SV2A density.** Correlation scatterplots between synaptophysin and SV2A density are shown for all cases (top left), controls (top right), PD (bottom left) and DLB (bottom right). Synaptophysin and SV2A density weakly correlated across all cases and in the control, PD and DLB group, while they did not correlate in the PDD group. Every data point represents one averaged measurement per layer III and VI, and it is color-coded by group (legend bottom right). **Legend**: *CTRL: control; DLB: Dementia with Lewy Bodies; PD: Parkinson's disease; PDD: Parkinson's disease dementia.*

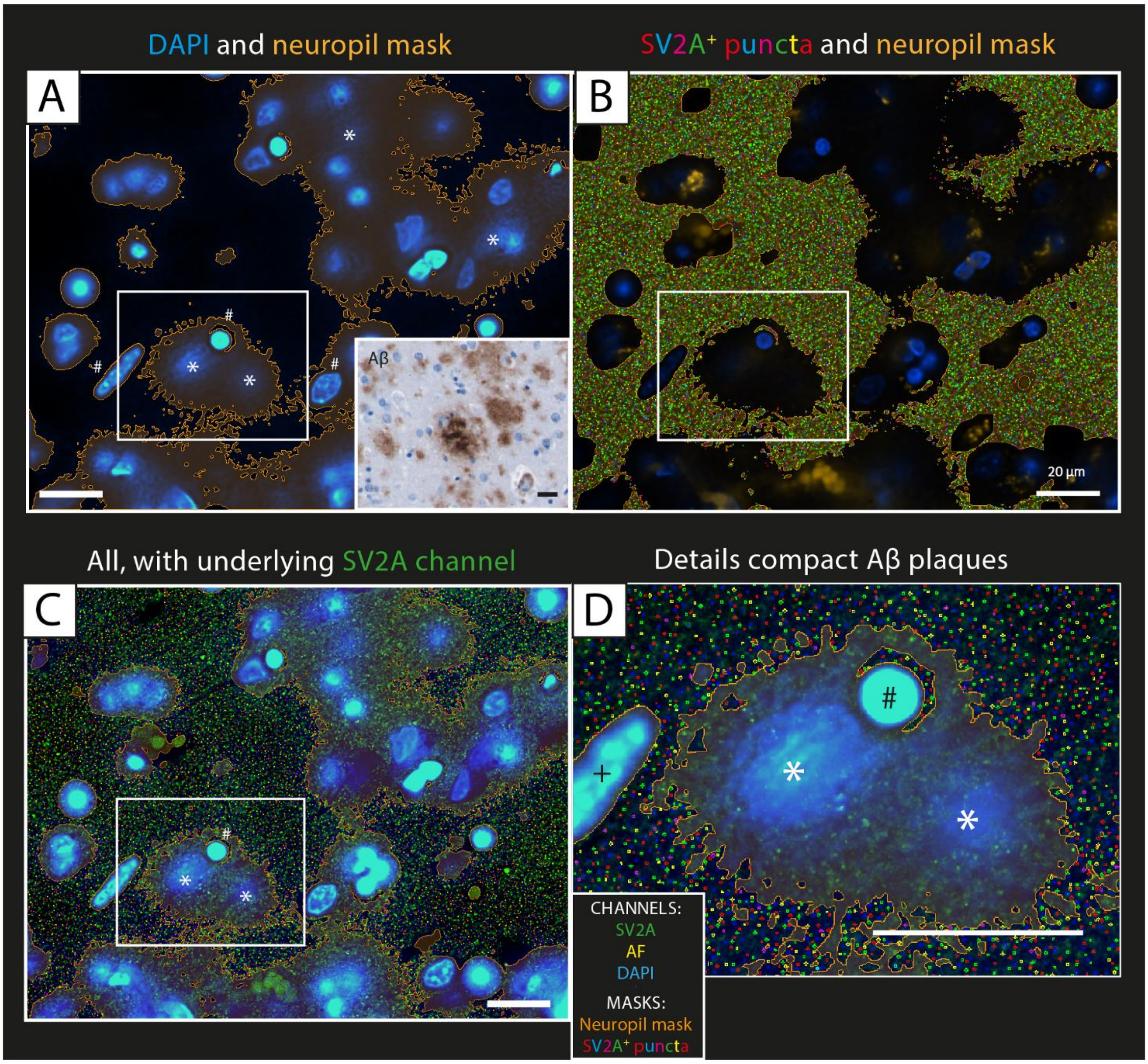


**Fig. S10 Relationship between amyloid-β and SV2A+ puncta: neuropil mask.** SV2A analysis of the posterior cingulate gyrus of a DLB case, layer III. In **(A)** we can see the DAPI channel (blue) together with the resulting neuropil mask (orange). From this image we can appreciate that what looks like amyloid-β (Aβ) compact plaques (*), positive to DAPI, besides the nuclei (#). In a consecutive section (insert of A), we can visualize amyloid-β compact plaques in the bright-field amyloid-β staining of the same case. **(B)** shows the next step of the analysis, that is the quantification of SV2A+ puncta (colorful dot-like mask) with the bright spot function only in the neuropil mask, therefore excluding not only the nuclei, the autofluorescence signal, and holes in the tissue, but also amyloid-β compact plaques. In **(C)** we can see the original staining, with DAPI (blue) and SV2A (green) with the masking explained in **(A)** and **(B). (D)** shows a zoom-in detail of two compact plaques (*) next to a nucleus (#) and a blood vessel (+). Particularly, here we can appreciate that the quantification of SV2A+ puncta (colorful dot-like mask) is carried out outside the nuclei and the compact plaques, while SV2A positive puncta (green dots) within the amyloid-β compact plaques are excluded from the analysis. The scale bar is 20 µm in all images. **Legend**: *Aβ: amyloid-β; AF: autofluorescence*.

### References supplementary figures


1. Adler DH, Pluta J, Kadivar S, Craige C, Gee JC, Avants BB, Yushkevich PA: **Histology-derived volumetric annotation of the human hippocampal subfields in postmortem MRI.** *Neuroimage* 2014, **84:**505-523.
2. Insausti R, Munoz-Lopez M, Insausti AM, Artacho-Perula E: **The Human Periallocortex: Layer Pattern in Presubiculum, Parasubiculum and Entorhinal Cortex. A Review.** *Front Neuroanat* 2017, **11:**84.
3. Bankhead P, Loughrey MB, Fernandez JA, Dombrowski Y, McArt DG, Dunne PD, McQuaid S, Gray RT, Murray LJ, Coleman HG, et al: **QuPath: Open source software for digital pathology image analysis.** *Sci Rep* 2017, **7:**16878.
4. Kovacs GG, Wagner U, Dumont B, Pikkarainen M, Osman AA, Streichenberger N, Leisser I, Verchère J, Baron T, Alafuzoff I: **An antibody with high reactivity for disease-associated α-synuclein reveals extensive brain pathology.** *Acta neuropathologica* 2012, **124:**37-50.
