## Supplementary tables for "Regional differences in synaptic degeneration are linked to alpha-synuclein burden and axonal damage in Parkinson’s disease and Dementia with Lewy bodies"

#### Table S1. Donor characteristics.

| *Clinical diagnosis* | Sex | Age at diagnosis (years) | Disease duration (years) | MDI (years) | CDR | Age at death (years) | PMD (hr:min) | Cause of death | ABC score [3] | Thal phase [4] | Braak NFT stage [1] | Braak α-syn stage [2] |
|---|---|---|---|---|---|---|---|---|---|---|---|---|
| *Ctrl* | M | - | - | - | n.a. | 68 | 8:30 | Euthanasia | A1B1C0 | 2 | 1 | 0 |
| *Ctrl* | F | - | - | - | n.a. | 63 | 8:10 | Euthanasia | A0B0C0 | 0 | 0 | 0 |
| *Ctrl* | M | - | - | - | n.a. | 82 | 10:40 | Liver cirrhosis | A1B1C0 | 1 | 1 | 0 |
| *Ctrl* | M | - | - | - | n.a. | 85 | 9:22 | Euthanasia | A1B1C0 | 1 | 1 | 0 |
| *Ctrl* | M | - | - | - | n.a. | 67 | 7:35 | Euthanasia | A1B1C0 | 1 | 2 | 0 |
| *Ctrl* | F | - | - | - | n.a. | 76 | 7:50 | Euthanasia | A1B1C0 | 2 | 1 | 0 |
| *Ctrl* | M | - | - | - | n.a. | 67 | 8:10 | Liver cirrhosis | A1B1C0 | 1 | 1 | 0 |
| *Ctrl* | F | - | - | - | n.a. | 77 | 9:00 | Euthanasia | A1B1C0 | 1 | 2 | 0 |
| *Ctrl* | M | - | - | - | n.a. | 57 | 9:50 | Euthanasia | A1 B0 C0 | 1 | 0 | 0 |
| *Ctrl* | F | - | - | - | n.a. | 83 | 5:50 | Euthanasia | A2 B2 C0 | 4 | 3 | 0 |
| *Ctrl* | F | - | - | - | n.a. | 87 | 8:35 | Urinary tract infection | A0B1C0 | 0 | 1 | 0 |
| *Ctrl* | F | - | - | - | n.a. | 72 | 7:20 | Heart failure | A0B0C0 | 0 | 0 | 0 |
| *Ctrl* | F | - | - | - | n.a. | 69 | 12:45 | Pulmonary embolism | A1B1C0 | 1 | 1 | 1 |
| *Ctrl* | M | - | - | - | n.a. | 59 | 8:00 | Euthanasia | A1B1C0 | 2 | 1 | 0 |
| *Ctrl* | M | - | - | - | n.a. | 77 | 11:25 | Pneumonia | A1B1C0 | 1 | 1 | 0 |
| *Ctrl* | F | - | - | - | n.a. | 78 | 10:00 | Unknown | A1B1C0 | 1 | 1 | 1 |
| *Ctrl* | F | - | - | - | n.a. | 77 | 4:35 | Unknown | A1B1C0 | 2 | 2 | 2 |
| *Ctrl* | F | - | - | - | n.a. | 59 | 8:10 | Euthanasia | A0B0C0 | 0 | 0 | 0 |
| *Ctrl* | F | - | - | - | n.a. | 71 | 6:50 | Lung carcinoma | A1B1C0 | 2 | 1 | 0 |
| *Ctrl* | M | - | - | - | n.a. | 74 | 10:20 | Euthanasia | A2B1C0 | 3 | 2 | 0 |

| *Clinical diagnosis* | Sex | Age at diagnosis (years) | Disease duration (years) | MDI (years) | CDR | Age at death (years) | PMD (hr:min) | Cause of death | ABC score [3] | Thal phase [4] | Braak NFT stage [1] | Braak α-syn stage [2] |
|---|---|---|---|---|---|---|---|---|---|---|---|---|
| *PD* | F | 61 | 22 | - | n.a. | 83 | 10:35 | Euthanasia | A0B1C0 | 0 | 1 | 5 |
| *PD* | F | 54 | 15 | - | 0.5 | 69 | 07:05 | Aspiration pneumonia | A1B1C0 | 2 | 2 | 6 |
| *PD* | F | 65 | 17 | - | n.a. | 82 | 09:17 | Aspiration pneumonia | A1B1C0 | 2 | 2 | 6 |
| *PD* | M | 61 | 17 | - | 0.5 | 78 | 07:15 | Euthanasia | A1B1C0 | 1 | 2 | 6 |
| *PD* | M | 76 | 16 | - | n.a. | 92 | 10:10 | Myocardial infarction | A2B2C1 | 3 | 3 | 4 |
| *PD* | M | 55 | 20 | - | 0.5 | 75 | 4:55 | End-stage PD | A2B1C0 | 3 | 2 | 6 |
| *PD* | M | 58 | 20 | - | n.a. | 78 | 3:30 | End-stage PD | A1B1C0 | 1 | 2 | 6 |
| *PD* | M | 70 | 23 | - | n.a. | 93 | 10:40 | Euthanasia | A1B2C0 | 2 | 4 | 6 |
| *PD* | M | n.a. | n.a. | - | n.a. | 83 | 6:30 | Euthanasia | A1B2C0 | 1 | 3 | 6 |
| *PD* | M | n.a. | n.a. | - | n.a. | 84 | 10:30 | Euthanasia | A1B2C0 | 1 | 3 | 6 |
| *PD* | M | 72 | 13 | - | 0.5 | 85 | 9:30 | End-stage PD | A1B1C0 | 1 | 2 | 5 |
| *PD* | M | 68 | 19 | - | 0.5 | 87 | 10:10 | Euthanasia | A2B1C0 | 3 | 2 | 6 |
| ***Clinical diagnosis*** | **Sex** | **Age at diagnosis (years)** | **Disease duration (years)** | **MDI (years)** | **CDR** | **Age at death (years)** | **PMD (hr:min)** | **Cause of death** | **ABC score [3]** | **Thal phase [4]** | **Braak NFT stage [1]** | **Braak α-syn stage [2]** |
| *PDD* | F | 67 | 16 | 11 | 3 | 83 | 10:40 | End-stage PDD | A3B2C2 | 4 | 4 | 6 |
| *PDD* | F | 84 | 10 | 7 | 3 | 94 | 06:50 | Femur fracture | A3B2C2 | 4 | 4 | 6 |
| *PDD* | F | 62 | 12 | 10 | 1 | 74 | 08:10 | Euthanasia | A2B1C0 | 3 | 2 | 6 |
| *PDD* | M | 58 | 21 | 16 | 2 | 79 | 09:25 | Subarachnoid bleeding | A2B1C1 | 3 | 2 | 6 |
| *PDD* | M | 44 | 18 | 2 | n.a. | 62 | 05:10 | End-stage PDD | A1B1C0 | 1 | 2 | 6 |
| *PDD* | M | 66 | 8 | 7 | 2 | 74 | 8:50 | Aspiration pneumonia | A1B2C0 | 2 | 3 | 6 |
| *PDD* | F | 62 | 19 | 16 | n.a. | 81 | 5:30 | End-stage PDD | A2B1C0 | 3 | 2 | 6 |
| *PDD* | M | 62 | 8 | 7 | 1 | 70 | 6:55 | Euthanasia | A1B1C0 | 1 | 1 | 6 |
| *PDD* | F | n.a. | n.a. | n.a. | n.a. | 74 | 9:30 | Euthanasia | A1B1C0 | 1 | 1 | 6 |
| ***Clinical diagnosis*** | **Sex** | **Age at diagnosis (years)** | **Disease duration (years)** | **MDI (years)** | **CDR** | **Age at death (years)** | **PMD (hr:min)** | **Cause of death** | **ABC score [3]** | **Thal phase [4]** | **Braak NFT stage [1]** | **Braak α-syn stage [2]** |
| *DLB* | M | 62 | 4 | 0 | 1 | 66 | 08:00 | Euthanasia | A3B2C2 | 4 | 4 | 6 |

| | | | | | | | | | | | | |
|---|---|---|---|---|---|---|---|---|---|---|---|---|
| *DLB* | M | 67 | 5 | 0 | 2 | 72 | 07:10 | Euthanasia | A2B1C2 | 3 | 2 | 6 |
| *DLB* | M | 70 | 7 | 0 | 3 | 77 | 07:15 | End-stage DLB | A1B2C0 | 1 | 3 | 6 |
| *DLB* | M | 88 | 3 | 0 | 2 | 91 | 4:30 | Euthanasia | A2B2C2 | 3 | 4 | 6 |
| *DLB* | F | 80 | 6 | 0 | 3 | 86 | 7:00 | Dehydration, cachexia | A3B2C2 | 5 | 4 | 6 |
| *DLB* | M | n.a. | n.a. | n.a. | n.a. | 61 | 10:10 | Unknown | A3B3C3 | 5 | 6 | 6 |

**Legend**: *α-syn: alpha-synuclein; Ctrl: control; DLB: Dementia with Lewy Bodies; F: female; hr: hour; IHC: immunohistochemistry; LB: Lewy body; M: male; min: minute; NFT: neurofibrillary tangles; PD: Parkinson's disease; PDD: Parkinson's disease dementia; PMD: post-mortem delay.*

**Table S2. Information on primary and secondary antibodies.**

| ***Antibody*** | **Antigen** | **Species** | **Origin details** | **Dilution** | **Incubation time** | **Antigen retrieval** | **Detection method** |
|---|---|---|---|---|---|---|---|
| ***Primary antibodies*** | | | | | | | |
| *pSer129 aSyn, clone EP1536Y* | Alpha synuclein phosphorylated at Ser129 | Rabbit igG | Abcam, Cambridge, UK | 1:8000 for BF | 4°C o.n. | Tris EDTA buffer (pH 9.0) in steam cooker | EnVision (HRP) |
| *Aβ, clone 4G8* | Aβ amino acid sequence 17-24 | Mouse igG2b | BioLegend, San Diego, USA | 1:8000 for BF | 4°C o.n. | Citrate buffer (pH 6.0) in steam cooker | EnVision (HRP) |
| *p-tau, clone AT8* | Tau phosphorylated at Ser202 and Thr205 | Mouse igG1 | ThermoFisher, Pittsburgh, USA | 1:800 for BF | 4°C o.n. | Citrate buffer (pH 6.0) in steam cooker | EnVision (HRP) |
| *Synaptophysin* | Recombinant protein of C-terminal cytoplasmic domain | Mouse igG1 | Agilent DAKO, Santa Clara, USA | 1:50 for FL | 4°C o.n. | Tris EDTA buffer (pH 9.0) in steam cooker | DoAM Alexa488 |
| *SV2A* | Clone EPR23500-32 | Rabbit igG | Abcam, Cambridge, UK | 1:50 for FL | 4°C o.n. | Tris EDTA buffer (pH 9.0) in steam cooker | DoAR Alexa488 |

| | | | | | | | |
|---|---|---|---|---|---|---|---|
| *NfL* | Immunogen corresponds to AA 1 to 284 (with AA 200-292 missing) | Rabbit igG | Synaptic systems, Göttingen, Germany | 1:600 for BF | 4°C o.n. | Tris EDTA buffer (pH 9.0) in steam cooker | EnVision (HRP) |
| ***Secondary antibodies*** | **Host species** | **Target species** | **Origin details** | **Dilution** | **Incubation time** | **Antigen retrieval** | **Detection method** |
| *DoAM Alexa 488* | Donkey IgG | Mouse | ThermoFisher, Pittsburgh, USA | 1:200 | 2 hrs at RT | / | fluorochrome |
| *DoAR Alexa 488* | Donkey IgG | Rabbit | ThermoFisher, Pittsburgh, USA | 1:200 | 2 hrs at RT | / | fluorochrome |

**Legend**: *BF: bright-field; DoAM: donkey anti-mouse; DoAR: donkey anti-rabbit; FL: fluorescence; hrs: hours; o.n.: overnight; RT: room temperature.*

**Table S3. Information on fluorescence 60x oil objective scanning at Olympus VS200.**

| **Channel** | **Target** | **Excitation wavelength** | **Filter** | **Exposure time** |
|---|---|---|---|---|
| **Synaptophysin staining** | | | | |
| DAPI | Nuclei | 378/52 nm | 432/36 nm | 1.5 ms |
| Alexa 488 | Synaptophysin | 474/27 nm | 515/36 nm | 50 ms |
| Autofluorescence | Autofluorescence | 474/27 nm | 595/31 nm | 50 ms |
| **SV2A staining** | | | | |
| DAPI | Nuclei | 378/52 nm | 432/36 nm | 1.5 ms |
| Alexa 488 | SV2A | 474/27 nm | 515/36 nm | 70 ms |
| Autofluorescence | Autofluorescence | 474/27 nm | 595/31 nm | 70 ms |

**Table S4. Contribution of all pathological markers to synaptic density in regions affected at Braak α-synuclein stage 4, 5, and 6.**

| Synaptophysin density | LB density | Amyloid-beta load | P-tau load | NfL immunoreactivity |
|---|---|---|---|---|
| **Braak 4** | β= -0.051, p=1.000 | β= -0.004, p=1.000 | β= 0.009, p=1.000 | *β= -0.009, p=0.125* |
| **Braak 5** | *β= -0.066, p=0.267* | β= 0.099, p=0.719 | β= -0.057, p=1.000 | *β= -0.008, p=0.230* |
| **Braak 6** | β= -0.038, p=1.000 | **β= 0.173, p=0.018** | *β= -0.156, p=0.245* | β= -0.005, p=0.405 |

| SV2A density | LB density | Amyloid-beta load | P-tau load | NfL immunoreactivity |
|---|---|---|---|---|
| **Braak 4** | **β= -0.095, p=0.008** | β= 0.057, p=1.000 | *β= 0.070, p=0.074* | *β= -0.006, p=0.525* |
| **Braak 5** | *β= -0.063, p=0.066* | **β= 0.194, p<0.001** | β= -0.014, p=1.000 | β= -0.007, p=0.239 |
| **Braak 6** | *β= -0.139, p=0.134* | **β= 0.332, p<0.001** | β= -0.024, p=1.000 | **β= -0.016, p<0.001** |

The tables show the contribution of all neuropathological markers to synaptophysin (top) and SV2A (bottom) density in regions affected at Braak α-synuclein stage 4, 5 and 6. Significant associations after multiple testing comparison correction are highlighted in **bold** (with red and blue colors indicating positive and negative associations, respectively), and associations that were significant before correction, but did not survive multiple testing comparison correction (Bonferroni method for 12 comparisons) are in *cursive* and underlined. We can see here that in areas affected at Braak stage 5, LB density was a negative predictor of both synaptophysin and SV2A density in these regions, however these associations did not survive multiple testing comparison correction.

**Legend:** *LB: Lewy body; NfL: neurofilament light chain.*

### References supplementary tables


1 Braak H, Alafuzoff I, Arzberger T, Kretzschmar H, Del Tredici K (2006) Staging of Alzheimer disease-associated neurofibrillary pathology using paraffin sections and immunocytochemistry. Acta neuropathologica 112: 389-404

2 Braak H, Del Tredici K, Rub U, de Vos RA, Jansen Steur EN, Braak E (2003) Staging of brain pathology related to sporadic Parkinson's disease. Neurobiol Aging 24: 197-211 Doi 10.1016/s0197-4580(02)00065-9

3 Montine TJ, Phelps CH, Beach TG, Bigio EH, Cairns NJ, Dickson DW, Duyckaerts C, Frosch MP, Masliah E, Mirra SS (2012) National Institute on Aging–Alzheimer's Association guidelines for the neuropathologic assessment of Alzheimer's disease: a practical approach. Acta neuropathologica 123: 1-11

4 Thal DR, Rüb U, Orantes M, Braak H (2002) Phases of Aβ-deposition in the human brain and its relevance for the development of AD. Neurology 58: 1791-1800
