## Supplementary results for "Regional differences in synaptic degeneration are linked to alpha-synuclein burden and axonal damage in Parkinson’s disease and Dementia with Lewy bodies"

### Age-related and sex differences in cortical synaptic density

Both synaptophysin and SV2A densities weakly decreased with age among all cases (synaptophysin: r = -0.06, *p=0.021*; SV2A: r = -0.10, *p=0.020*). Synaptophysin density was significantly higher in females compared to males across all cases (F/M: 20/27, +5%, *p<0.001*), especially in controls (F/M: 11/9, +7%, *p<0.001*), but not in PD (F/M: 3/9, *p=0.817*) nor PDD (F/M: 5/4, *p=0.096*) (**Fig. S5A**). The analysis was not performed in DLB cases since the female/male ratio was 5/1. SV2A density did not differ in females and males across all cases (*p=0.238*), which was the same for controls (*p=0.347*) and PD cases (*p=0.364*), while PDD cases showed higher SV2A density in males compared to females (F/M: 5/4, +5%, *p=0.049*) (**Fig. S5B**).

In summary, we found that both synaptophysin and SV2A density decreased with age, while only synaptophysin showed a sex-effect, being more abundant in females compared to males in controls.

### Cortical layer III and V-VI differences in synaptic density

Both synaptophysin and SV2A densities were slightly lower in layer V-VI compared to layer III across all cases and regions (synaptophysin: -3%, *p=0.006*; SV2A: -3%, *p=0.022*) **(Fig. S6)**. Particularly, synaptophysin density was lower in layer V-VI compared to layer III in PD (-4%, *p=0.043*) and controls (-4%, *p=0.039*), but this difference was lost in PDD (*p=0.419*) and DLB (*p=0.603*) (**Fig. S6C**). SV2A density did not show any group-specific layer difference (*p>0.05*) (**Fig. S6D**).

### Differential within-group regional distribution of synaptophysin and SV2A puncta

Synaptophysin density showed a similar distribution across brain areas in all groups (all p>0.05) **(Fig. S8 A,B)**. In turn, SV2A density showed different patterns across brain areas within groups **(Fig. S8 C,D).** In the control group, SV2A density was significantly lower in the middle temporal gyrus compared to the fusiform gyrus (-15%, p=0.024). In the PD group, SV2A density was significantly lower in the middle temporal gyrus compared to the entorhinal cortex (-12%, $p=0.028$) and the superior frontal gyrus (-12%, $p=0.032$). In the PDD group, almost all regions showed a lower SV2A density compared to the superior frontal gyrus and the posterior cingulate cortex (ENTC vs GFS: -13%, $p=0.043$; PHG vs GFS: -18%, $p<0.001$; FusG vs GFS: -18%, $p<0.001$; ACC vs GFS: -20%, $p<0.001$; MTG vs GFS: -17%, $p=0.001$; PHG vs PCC: -13%, $p=0.031$; FusG vs PCC: -14%, $p=0.019$; ACC vs PCC: -16%, $p=0.002$). Lastly in the DLB group, SV2A density was significantly lower in the anterior cingulate cortex compared to the superior frontal gyrus (-21%, $p=0.016$) and the posterior cingulate cortex (-20%, $p=0.029$). Overall, regions affected at later Braak stages (Braak stage 6, i.e. superior frontal and posterior cingulate cortex) seemed to be less affected in PD, PDD and DLB compared to regions affected at earlier stages.

As synaptophysin and SV2A density showed a different distribution patterns across brain regions, we investigated whether they correlated with each other. Overall, synaptophysin and SV2A density correlated weakly ($r = 0.20$, $p<0.001$). Specifically, they weakly correlated in PD ($r = 0.20$, $p=0.001$), DLB ($r = 0.28$, $p=0.007$), and controls ($r = 0.13$, $p=0.026$), but not in PDD ($p=0.151$) **(Fig. S9).**

In summary, synaptophysin had a similar density across brain regions, whereas SV2A density was more abundant in regions affected at Braak stage 6 compared to regions affected at Braak stage 4 and 5. Synaptophysin and SV2A densities only weakly correlated with each other.
